## Supplementary material for "Genomic screening of 16 UK native bat species through conservationist networks uncovers coronaviruses with zoonotic potential": Suppl. files: Supplementary Figure and Data legends 030523.docx

Tan et al.

**Supplementary Figure 1. RT-PCR assays underestimate coronavirus prevalence.** Heatmap summarising the number of mismatches of the forward (F) and reverse (R) degenerate primers described in previous studies to (a) novel genomes, and (b) to the nine novel and 2118 genomes in our custom coronavirus database. Both heatmaps are matched to the tips of the alignment-free trees generated from the genomes analysed, which are similar to that shown in Fig. 1a but represented as a linear phylogram. Heatmap cells coloured white or gray indicate no detectable homology between a degenerate primer and a genome by BLASTn.

**Supplementary Figure 2. Collection of faecal samples from 16 UK bat species through extensive network of bat rehabilitators.** (a) Temporal distribution of samples collected with the number of samples per host species annotated. (b) Geographical distribution of samples collected relative to the major cities in the UK.

**Supplementary Figure 3. Analysis of the UK bat faecal virome.** (a) Heatmap summarizing the number of samples per UK bat species where a particular viral family was present, based on Kraken2 taxonomic assignment of reads. Viral families that are known to infect mammals are highlighted in brown. (b) The total relative abundance of mammalian or non-mammalian viral species in each sample. Data are visualized with both Gaussian kernel probability density and box-and-whisker plots (centre line, median; box limits, upper and lower quartiles; whiskers, 1.5x interquartile range). A two-sided Mann-Whitney U test was used to test if the two distributions differed.

**Supplementary Figure 4. Even read coverage across all complete genomes recovered from UK bats.** Sequencing reads were mapped back to the final genomes using Bowtie2 and per-position read coverage was calculated using Samtools.

**Supplementary Figure 5. Genome schematics of the novel UK bat coronaviruses.** To-scale layouts of ORFs within the novel bat coronaviruses from this study compared to prototypic genomes from the same subgenera. ORF1ab polyproteins are shown in red, structural proteins in orange, accessory proteins in yellow, and putative novel ORFs in blue. Missing ORFs relative to the prototypes shown by dotted lines. Standard coronavirus gene nomenclature was used throughout. This figure was made using Adobe Illustrator v27.1.1 and Geneious v11.1.5 ([https://www.geneious.com](http://www.geneious.com/)).

**Supplementary Figure 6. Species distribution maps of UK bats.** (a) Predicted distributions of *R. ferrumequinum* and *R. hipposideros* species in the UK. (b) Species diversity (i.e., number of species) found within a 5x5 km square grid computed based on occurrence records dating from 2000-present. (c) Predicted species diversity all 17 UK breeding bat species found within a 1x1 km square grid. All predicted distributions were generated by our ensemble machine learning model. Species were deemed to be present if the predicted probability score (i.e., habitat suitability) generated for any square grid exceeds 0.8. *Rhinolophus* samples and all UK bat samples where coronavirus genomes or partial contigs were recovered, and whose exact geographical coordinates were available are annotated in (a) and (c), respectively.

**Supplementary Figure 7. Western blot analyses of spike pseudoviruses and cell receptor expression.** (a) Western blot showing relative ACE2 expressions of stably transduced, transfected or non-transfected/transduced HEK293T. (b) Western blot analysis of HEK293T cells transfected with different ACE2 constructs. All ACE2 proteins tagged with C-terminal HA tag. Equal loading shown by probing with anti-tubulin antibody. (c) Western blot analysis of concentrated pseudovirus expressing different sarbecovirus, merbecovirus and pedacovirus spike proteins. Sarbecovirus spike expression (upper panel) determined by a pan-sarbecovirus anti-S2 antibody. Pedacovirus and merbecovirus spike expression determined by incorporatation of C-terminally Myc-tagged spike (lower panel). The upper band corresponds to uncleaved, full length spike, the lower band to the cleaved S2 fragment. Loading shown by p24 lentiviral capsid protein. All western blots shown are representative repeats of n=3 independent experiments performed.

**Supplementary Figure 8. Protein surfaces of hACE2 in contact with RhGB07 or SARS-CoV-2 receptor-binding domain (RBD).** The structure of hACE2 is shown in grey and the surface in contact with the RBDs of RhGB07 (blue) and SARS-CoV-2 (orange) are highlighted. We computed the surface are of hACE2 in contact with either RhGB07 or SARS-CoV-2 RBD using the *buriedarea* command in *ChimeraX.*

**Supplementary Figure 9. European sarbecoviruses posses an RAKQ motif resembling a furin cleavage site.** (a) Sequence alignment of sarbecovirus spike genes at the region surrounding the SARS-CoV-2 furin cleavage site (FCS) and R-A-K-Q furin cleavage site precursor in UK sarbecoviruses. Sequence alignment was visualized using UGENE v42.0. The alignment region comprising SARS-CoV-2 spike residue positions 667-699 is indicated by a black rectangle and corresponds to the extended S1/S2 loop containing the R-R-A-R FCS present in SARS-CoV-2. Barchart showing the proportion of genomes with residues identical to SARS-CoV-2 at each position (top). Maximum-likelihood tree identical to that shown in Fig. 3c (left) showing the genetic relatedness of Asian, European and African sarbecoviruses. (b) Western blot of RhGB07 spike with or without the Q672R mutation (generating an RAKR motif). SARS-CoV-2 spike with or without the 678-NSPRRARS-687 deletion were used as negative and positive controls, respectively.

**Supplementary Figure 10. High prevalence of recombination amongst sarbecoviruses.** (a) Distribution of recombination events detected by at least six of the nine recombination detection algorithms in RDP4. This analysis was performed on an alignment of 218 representative sarbecoviruses, including RhGB01 and our four novel sarbecoviruses (RhGB07, RhGB08, RfGB01, RfGB02), using NC_025217 as the reference. (b) All recombination events involving RhGB01-like viruses either as donor or recipients. Recombination events were supported by 2-6 detection algorithms.

**Supplementary Figure 11. Species distribution modelling for the 17 UK breeding bat species.** (Left) Performance of individual machine-learning algorithms in predicting species distributions. (Right) Maps of individual species distributions. Predicted probability scores indicate the predicted habitat suitability for each 1x1km square grid, which ranges from 0 (unsuitable habitat) to 1 (suitable habitat). The number of occurrence records for each bat species used to train the models, and the geographical locations of bat samples collected in this study are indicated.

**Supplementary Figure 12. Raw uncropped images of western blots.** Panels (a), (b), (c) and (d) correspond to the images shown in Supplementary Fig. 7a, 7b, 7c and 9b, respectively.

**Supplementary Data 1.** Summary of a selection of coronavirus surveillance studies in bats.

**Supplementary Data 2.** Amino acid changes of RfGB02 from RhGB08.

**Supplementary Data 3.** Metadata of all samples sequenced by deep RNA sequencing.

**Supplementary Data 4.** Metadata of all 2118 coronavirus genomes from NCBI and GISAID used in this study.

**Supplementary Data 5.** List of NBN Atlas datasets used in this study.

**Supplementary Data 6.** GISAID acknowledgement table for GISAID genomes used in this study.
