## Supplementary material for "Genomic screening of 16 UK native bat species through conservationist networks uncovers coronaviruses with zoonotic potential": Suppl. files: Supplementary Information_030523.docx

Tan et al.

### Existing RT-PCR assays underestimate coronavirus prevalence in bats

Given that RT-PCR has conventionally been used to screen for coronaviruses in bats, we sought to determine if the novel coronavirus genomes we recovered from our metatranscriptomes could have been detected using published pan-coronavirus primers. Using BLASTn searches, we aligned the external RT-PCR primers that have been described previously^1–5^ against all coronavirus genomes in our custom database and our nine novel genomes. These include primers that have been used widely^1,3,4^, and updated primers described in two more recent studies^2,5^. Amongst these primers, the ones designed by Holbrook et al.^5^ are an updated version of those by Watanabe et al.^3^. Notably, whether a primer can bind to a particular genomic sequence is difficult to predict *in vitro* since the impact of mismatches on primer binding can depend on various factors such as the position of the mismatch or annealing temperature^6–8^. We therefore assumed that a primer sequence can bind to a coronavirus genome if a primer-genome alignment could be produced by BLASTn, and conversely, that a primer sequence is not likely to bind if no primer-genome alignment could be identified. Under this assumption, the coronavirus diversity that can be ‘detected’ by each primer set can be estimated by the proportion of coronavirus genomes that could be aligned to a query primer sequence. Since most of these primers contained degenerate bases, we performed the BLASTn analysis on every combination of non-degenerate bases for each primer and retained only the primer-genome alignment with the lowest number of mismatches.

None of the external primer sets, except that by Vijgen et al.^4^, were able to detect all nine novel genomes (Supplementary Figure 1a). In fact, three of the external primer sets^1–3^ could detect at most one of the novel coronaviruses. We extended this analysis further by analysing the sequence homology of all external primer sets to all genomes in our custom coronavirus database. All external primer sets carried at least one mismatch or had no detectable homology to at least one coronavirus genome in our database, indicating that none are likely to capture the full existing diversity of coronaviruses (Supplementary Figure 1b). Strikingly, the proportion of coronavirus genomes that could be detected by any external primer set, estimated from the number of detectable primer-genome alignments, ranged from 9.5 to 93.5%. Given that our analysis only includes the external primers, additional mismatches in the internal primer set may exacerbate the poor sensitivity of these RT-PCR assays. Overall, these findings indicate that RT-PCR screens that employ these primers likely underestimate viral prevalence in the systems being studied.

### Genome structure analyses indicate the presence of novel genes

We used various bioinformatic tools (see Methods) to determine if these genomes carry any novel genes. No notable novel genes were identified in the sarbecoviruses, which like RhGB01 have a similar genome structure to SARS-CoV-2 and SARS-CoV but are missing ORF8^9^ (Supplementary Figure 5). Although RfGB02 has an out of frame deletion that likely results in a truncated ORF7a. Similarly PpiGB02, MdGB02 and MdGB03 had similar genome structures to other bat Pedacoviruses, potentially expressing an additional ORF7 relative to PEDV^10^. The pedacovirus MdGB01 does however contain an additional potential ORF8 at the 3’ end of the genome, which is absent in the other UK bat pedacoviruses. This potential ORF8 has an upstream putative transcriptional regulatory sequence (TRS) and would result in expression of a 56 amino acid (a.a.) protein. However, PaGB01 encodes a novel 100 a.a protein that is only 54.9% similar to its closest homologue, the ORF3 accessory protein in MERS-CoV. This putative ORF3-like protein could not be assigned to any InterPro protein families^11^, but was predicted to contain a transmembrane and an extracellular domain. PaGB01 also encodes a 218 a.a. protein at 73.3% identity to the MERS-CoV ORF5 protein. Finally PaGB01 also encodes an ORF predicted to express an 83 a.a. protein, partially overlapping (in the +1 reading frame) with its N gene at the 3’ end of its genome. Consistent with coronavirus gene naming conventions, this would be named ORF8c. The divergence of these novel proteins from MERS-CoV are largely in line with that between the accessory proteins in MERS-CoV and other bat-borne MERS-CoV-related species, btCoV-HKU4 and btCoV-HKU5^12^. This indicates that the novel proteins may possess functions similar to the MERS-CoV accessory proteins.

Accessory proteins are non-essential for coronavirus replication *in vitro*, but are thought to play key roles in host-virus interactions. For example, ORF3 and ORF5 proteins in MERS-CoV have been shown to induce apoptosis^13^ and also to antagonise interferon responses^14^, which are a key aspect of the innate immune response to viruses in humans. The accessory genes of coronaviruses are highly variable in number and function across the family *Coronaviridae*. However, MERS-CoV and its close bat-borne relatives, btCoV-HKU4 and btCoV-HKU5, share a similar number of accessory genes with similar functions, despite low protein sequence similarities between the accessory proteins from these species^12,15^. In light of this, further characterisation of the novel proteins identified in PaGB01 may reveal fundamental insights on the evolution of viral pathogenicity. For example, if the accessory genes of PaGB01 match the function of the MERS-CoV equivalent proteins in interacting with human cellular signalling pathways, that could suggest that immunoregulation is a conserved function amongst MERS-CoV-related coronaviruses and may help explain how MERS-CoV is able to cause human disease. Conversely, a lack of shared activity may indicate that these functions are unique to MERS-CoV and its closest relatives and are not universally found in other sister lineages, perhaps explaining why there is no evidence of other MERS-related virus infections in humans to date.

### High prevalence of recombination amongst sarbecoviruses

Given that further adaptations are necessary for the zoonotic emergence of RhGB01-like viruses, we asked if genetic recombination may speed up this process. Recombination in viruses allows the genetic transfer of large sections of the genome in a single event, helping them sample the genomic sequence space at a more rapid pace when compared to the accumulation of point mutations alone^16^. In fact several regions in the spike protein of coronaviruses that influence host range have been suggested to have been acquired through recombination^17^, which implies that recombination may be an important driver for zoonotic emergence. As such, we performed recombination analyses for sarbecoviruses, including our novel sequences, using the recombination detection program (RDP)^18^. This tool comprises a suite of algorithms for recombination detection and has been used previously for sarbecoviruses^19,20^. We searched for recombination amongst 218 representative sarbecovirus genomes using all nine algorithms implemented within RDP4 (RDP^21^, GENECONV^22^, BOOTSCAN^23^, MaxChi^24^, Chimaera^25^, SisScan^26^, PhylPro^27^, LARD^28^ and 3SEQ^29^), retaining predicted breakpoints supported by at least six of these methods. Using this approach, we detected 202 putative recombination events amongst the sarbecoviruses considered, suggesting a high prevalence of recombination within the subgenus. Additionally, we detect an overrepresentation of recombination signals near the N-terminal half of the spike protein (Supplementary Figure 11a), which also contains the receptor binding domain that is the primary determinant of host receptor usage. We also identified six recombination events within the RhGB01-like viruses supported by 2-6 detection algorithms (Supplementary Figure 11b), demonstrating the potential for recombination involving the novel UK sarbecoviruses. Overall, these results support frequent events of recombination in sarbecoviruses, which may increase the likelihood of novel sarbecoviruses, some which may be zoonotic, emerging in *Rhinolophus* bats in the UK.
