## Supplementary material for "Genomic screening of 16 UK native bat species through conservationist networks uncovers coronaviruses with zoonotic potential": Suppl. files: Supplementary Table 6 gisaid acknowledgement table.pdf

### SUPPLEMENTAL TABLE

**Data Availability**

GISAID Identifier: EPI_SET_230109gu
doi: 10.55876/gis8.230109gu

All genome sequences and associated metadata in this dataset are published in GISAID's EpiCoV database. To view the contributors of each individual sequence with details such as accession number, Virus name, Collection date, Originating Lab and Submitting Lab and the list of Authors, visit 10.55876/gis8.230109gu

**Data Snapshot**

- EPI_SET_230109gu is composed of 29 individual genome sequences.
- The collection dates range from 2010-12-06 to 2022-01-03;
- Data were collected in 4 countries and territories;
- All sequences in this dataset are compared relative to hCoV-19/Wuhan/WIV04/2019 (WIV04), the official reference sequence employed by GISAID (EPI_ISL_402124). Learn more at https://gisaid.org/WIV04.
