## Supplementary material for "Genomic screening of 16 UK native bat species through conservationist networks uncovers coronaviruses with zoonotic potential": Suppl. files: Supplementary Figure 11.pdf

***Barbestella barbastellus* (No. of occurrence records = 270)**

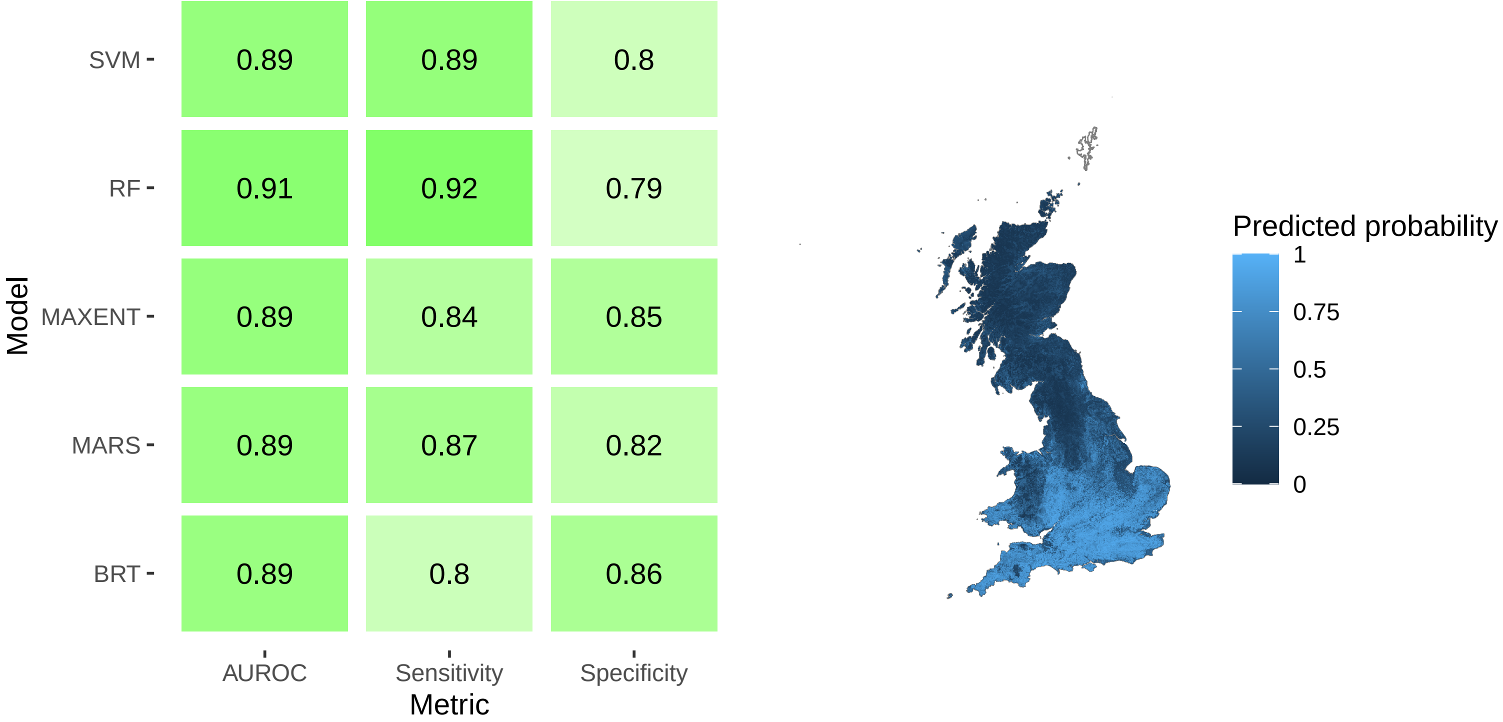

*Myotis alcathoe* (No. of occurrence records = 68)

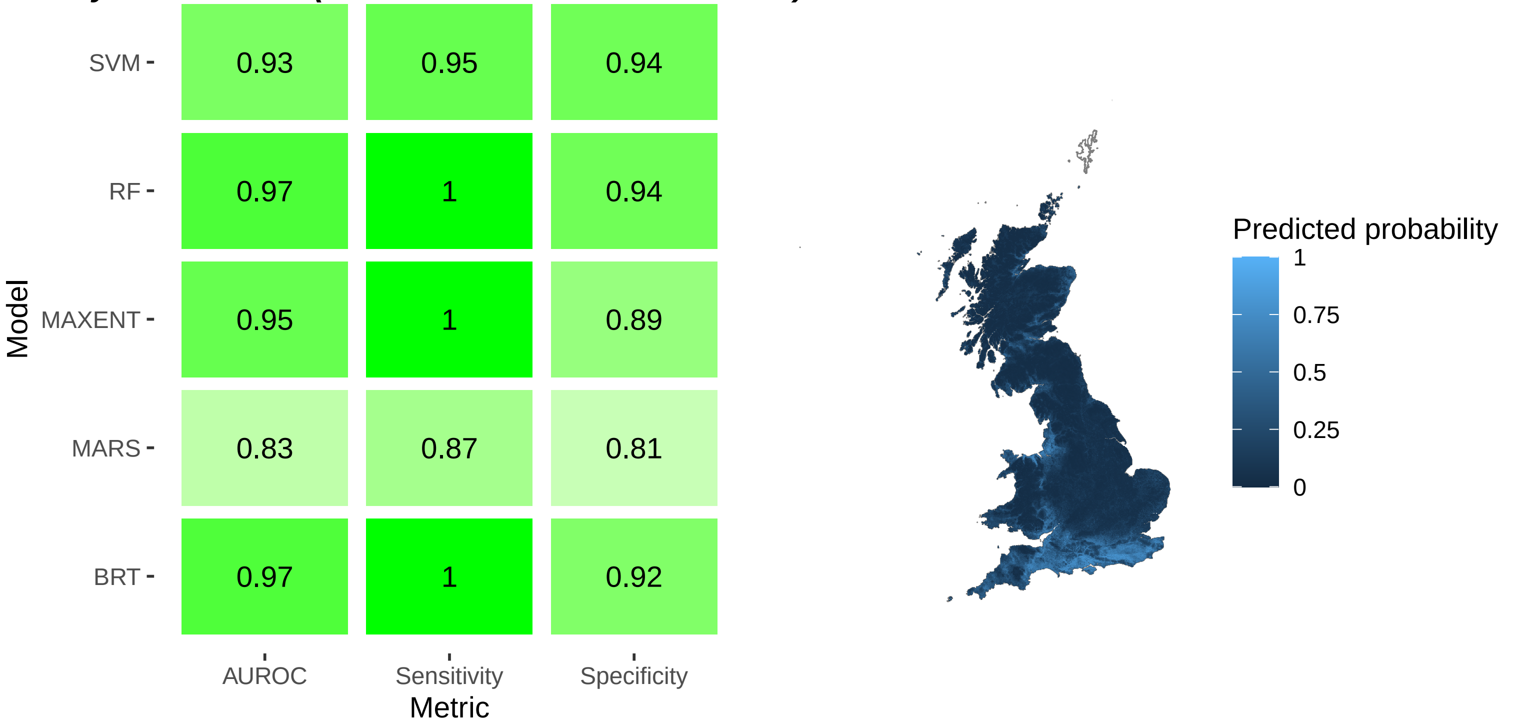

*Myotis bechsteinii* (No. of occurrence records = 73)

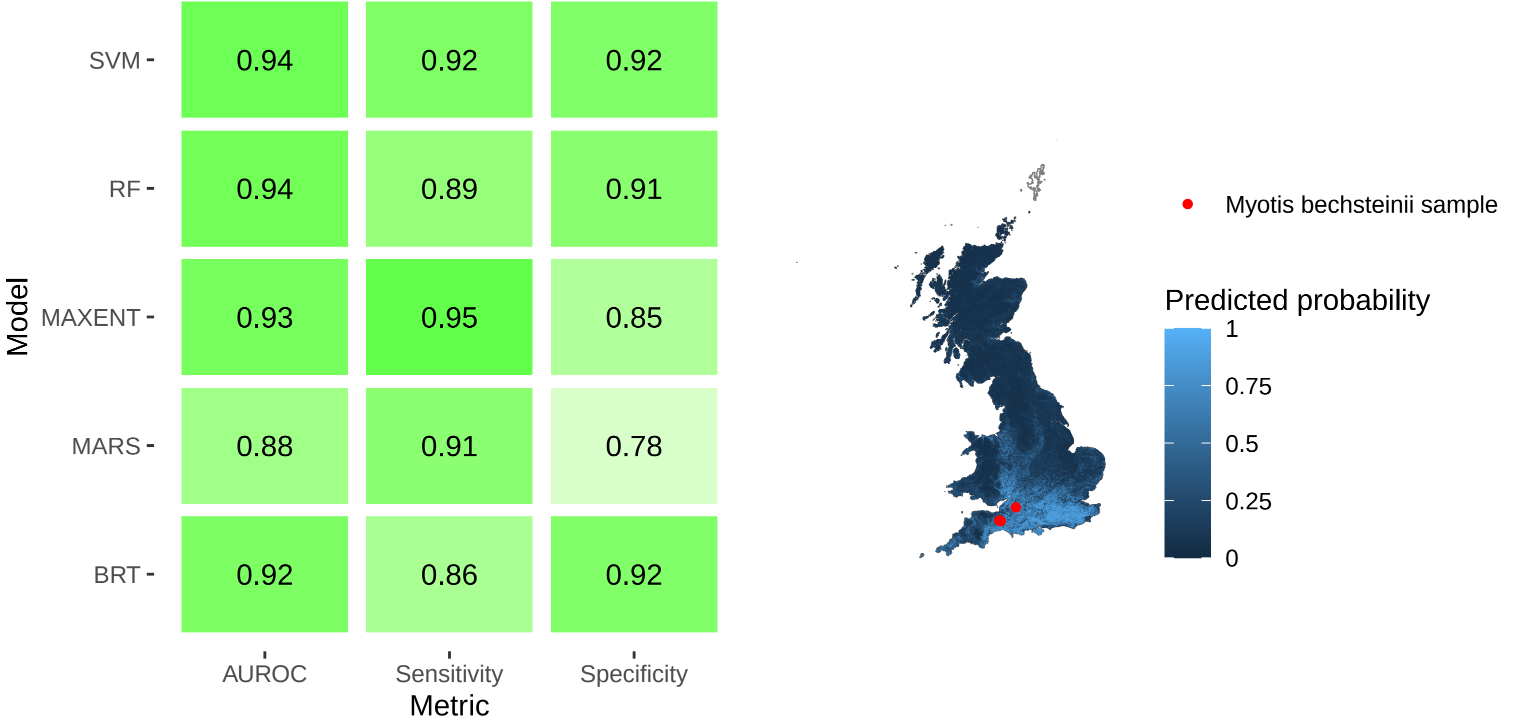

*Myotis brandtii (No. of occurrence records = 117)*

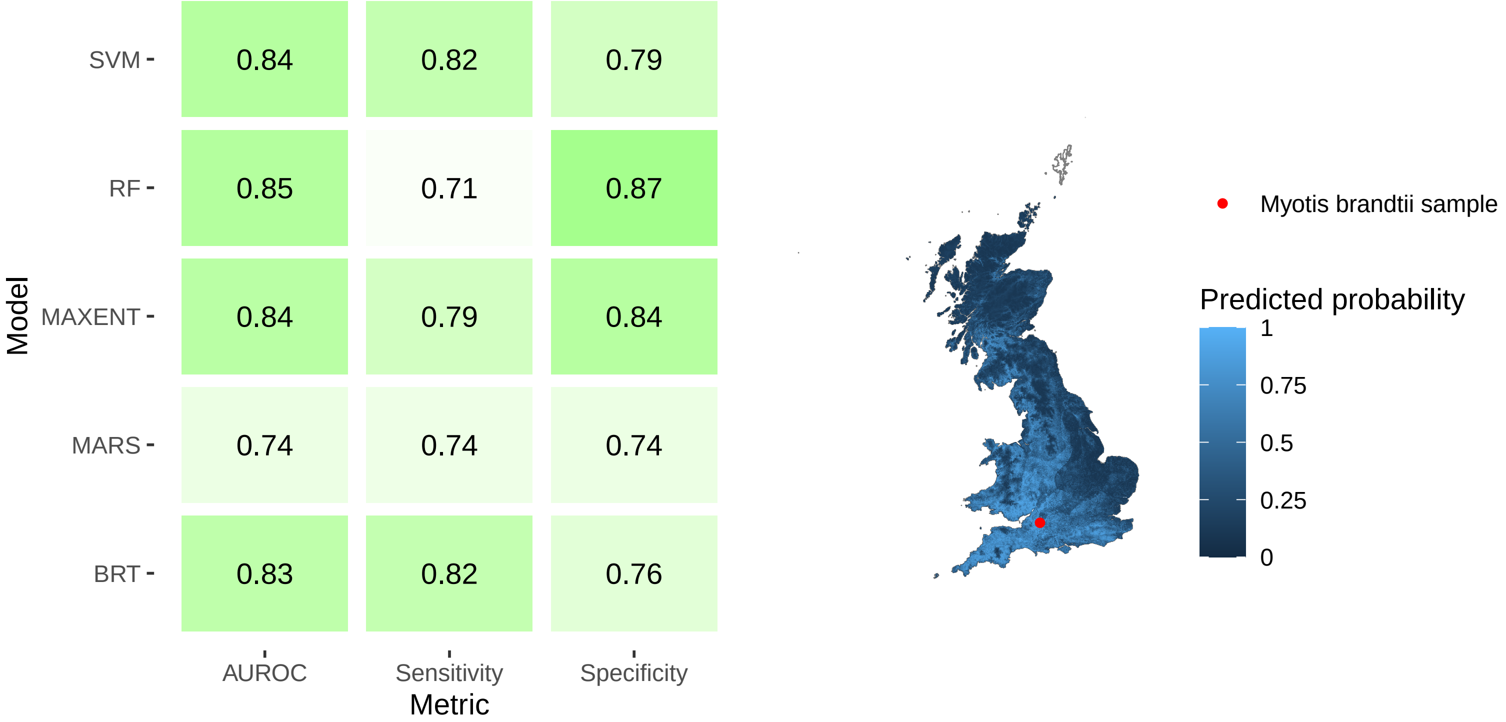

**_Plecotus auritus (No. of occurrence records = 4297)_**

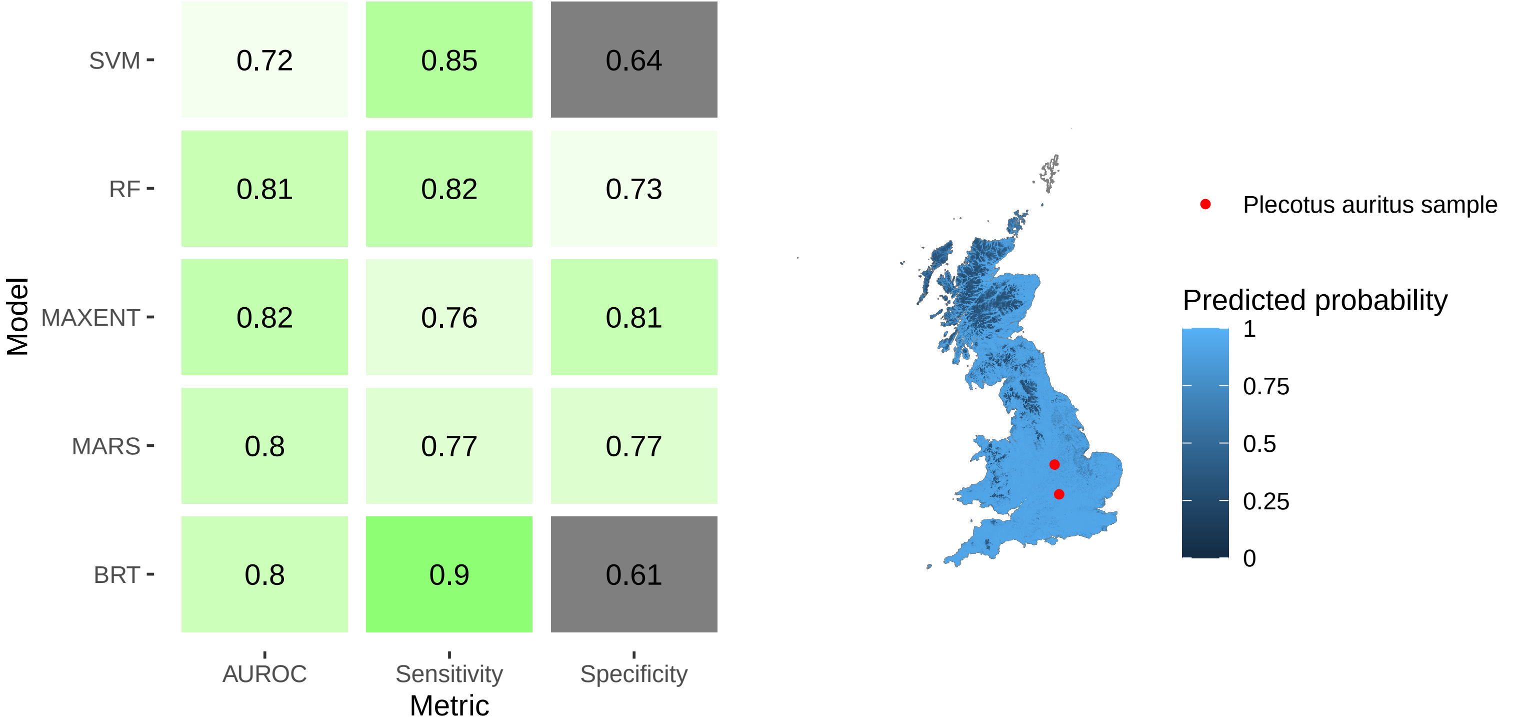

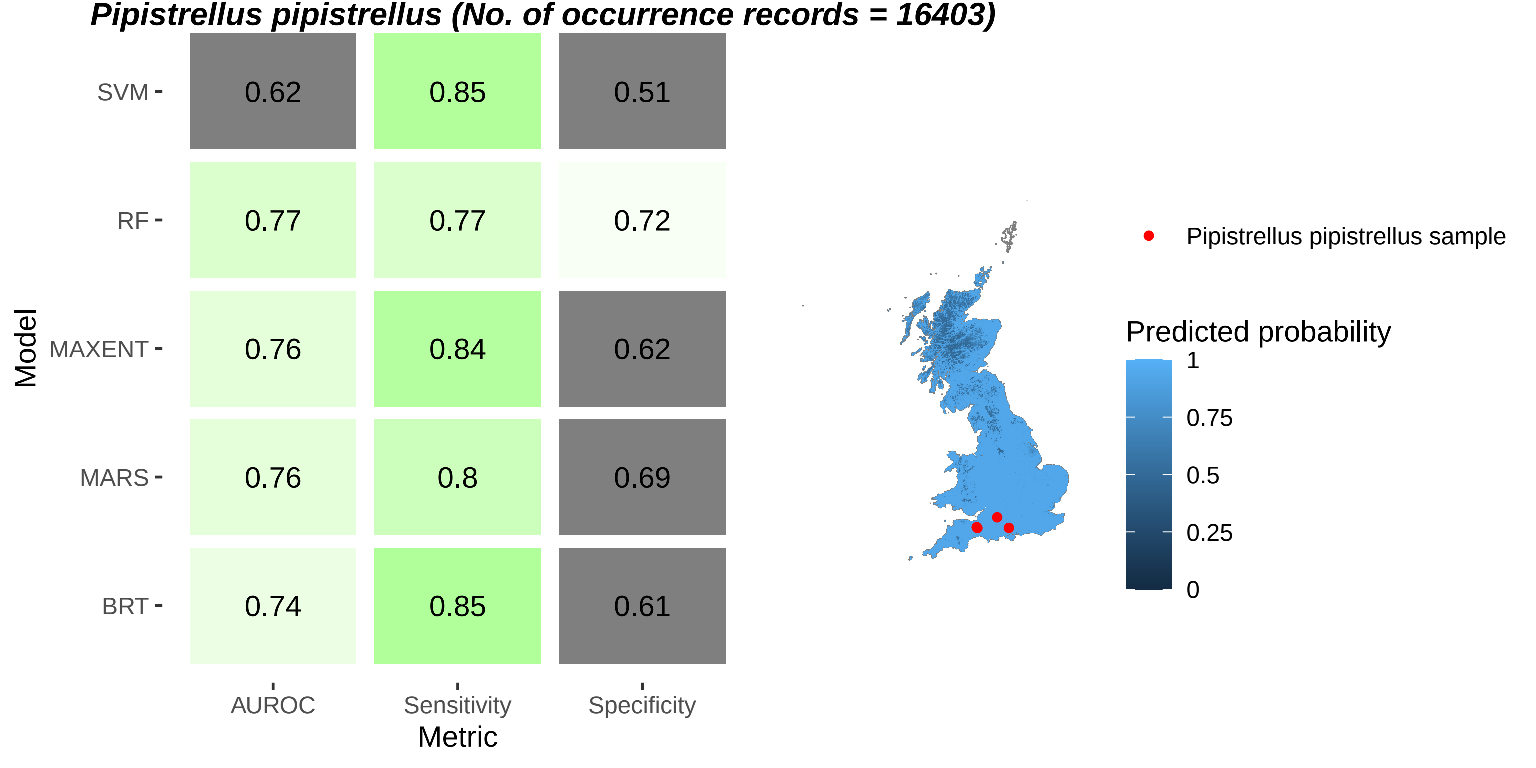
Pipistrellus pipistrellus (No. of occurrence records = 16403)
Model
SVM
RF
MAXENT
MARS
BRT
0.62
0.85
0.51
0.77
0.77
0.72
0.76
0.84
0.62
0.76
0.8
0.69
0.74
0.85
0.61
AUROC
Sensitivity
Specificity
Metric
Pipistrellus pipistrellus sample
Predicted probability
1
0.75
0.5
0.25
0

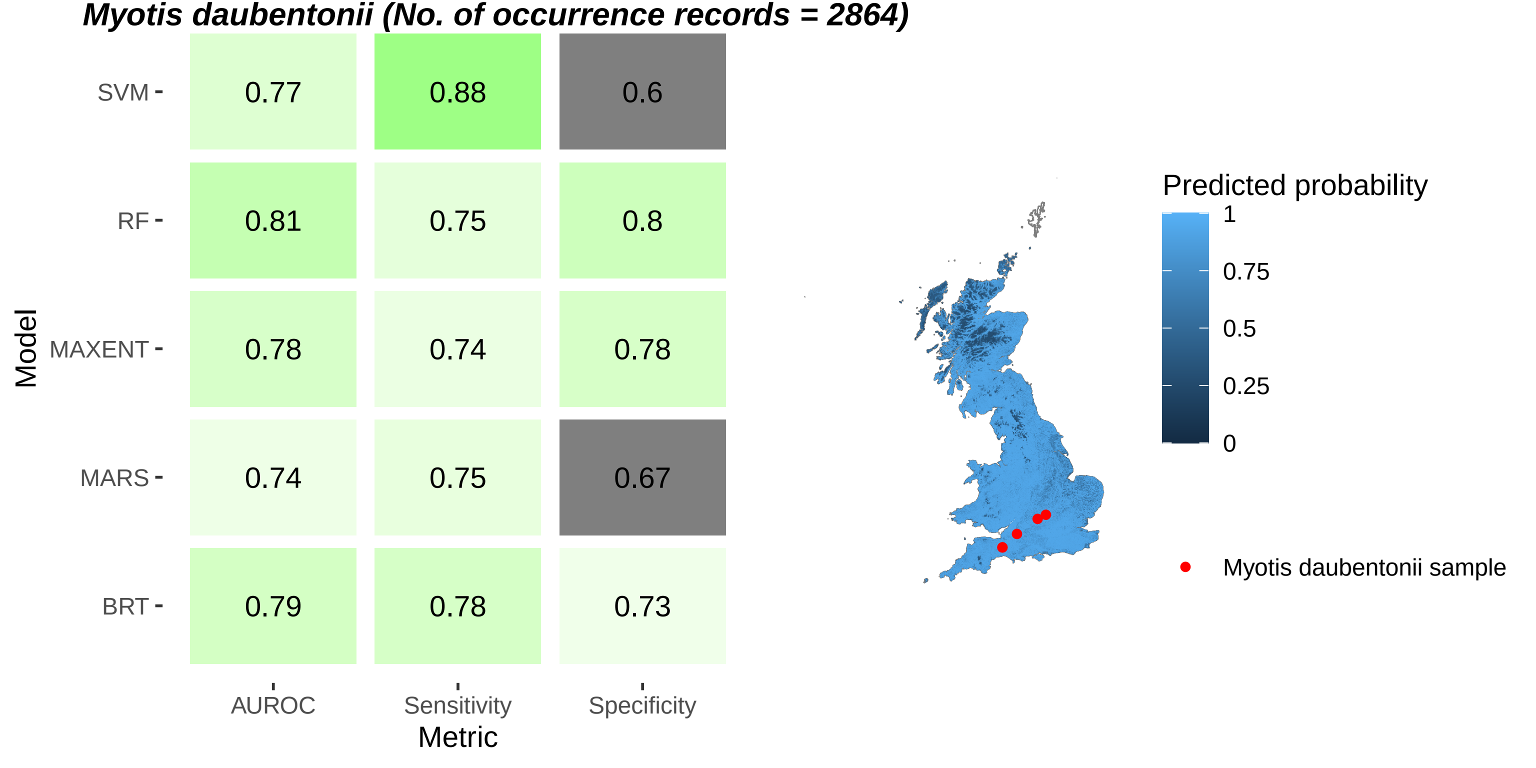
Myotis daubentonii (No. of occurrence records = 2864)
SVM
RF
MAXENT
MARS
BRT
Model
0.77
0.88
0.6
0.81
0.75
0.8
0.78
0.74
0.78
0.74
0.75
0.67
0.79
0.78
0.73
AUROC
Sensitivity
Specificity
Metric
Predicted probability
1
0.75
0.5
0.25
0
Myotis daubentonii sample

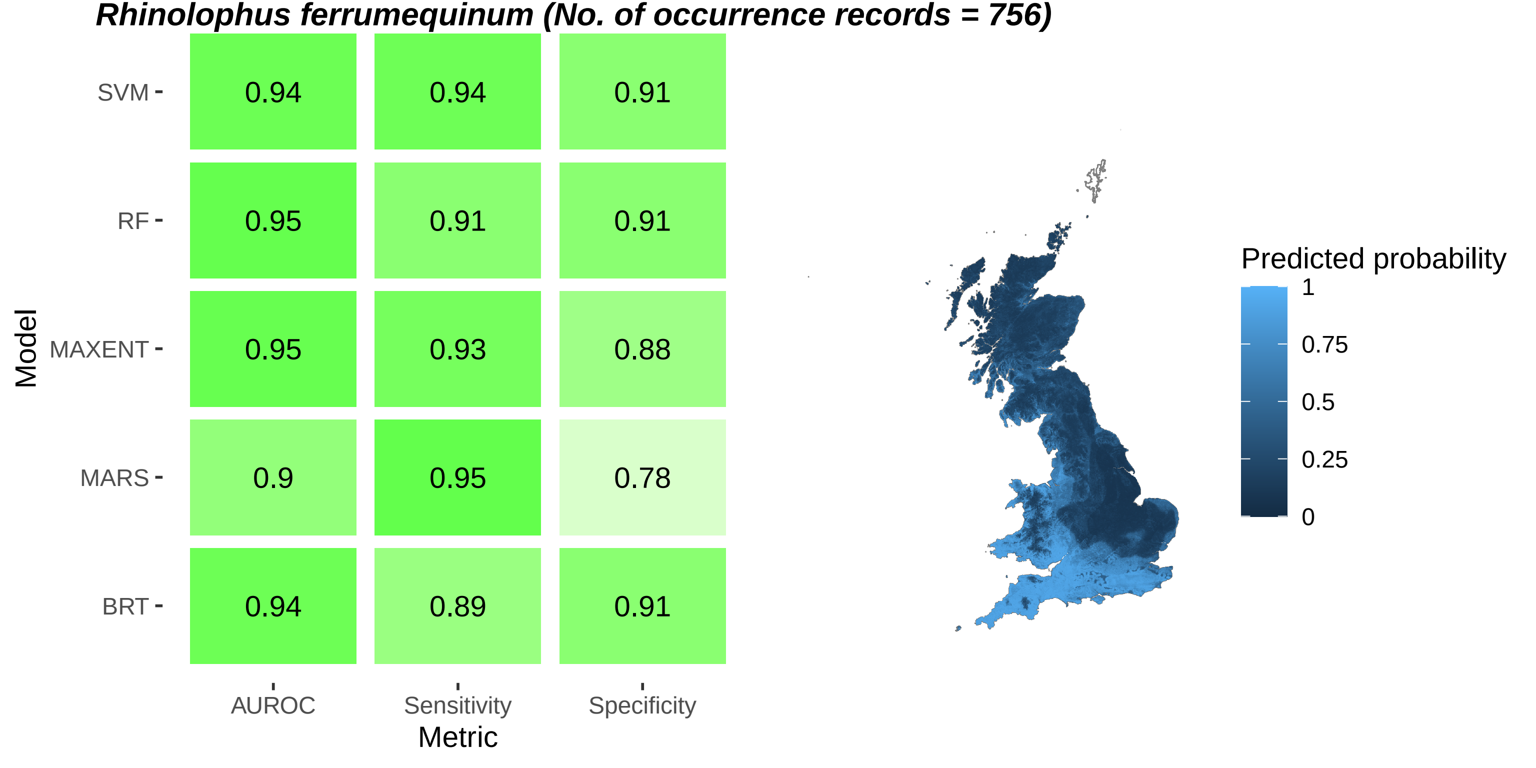
Rhinolophus ferrumequinum (No. of occurrence records = 756)
Model
SVM
RF
MAXENT
MARS
BRT
0.94
0.94
0.91
0.95
0.91
0.91
0.95
0.93
0.88
0.9
0.95
0.78
0.94
0.89
0.91
AUROC
Sensitivity
Specificity
Metric
Predicted probability
1
0.75
0.5
0.25
0

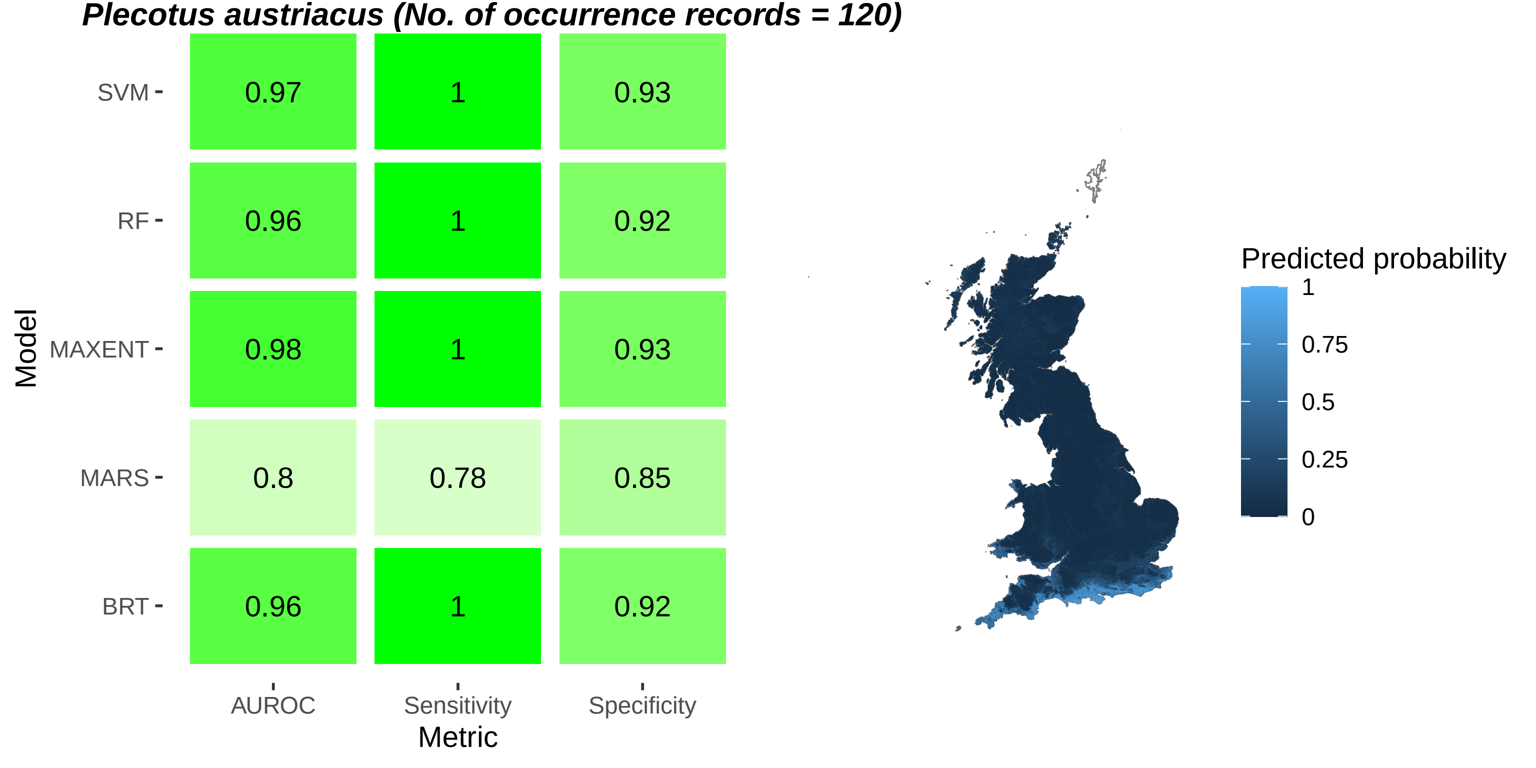
Plecotus austriacus (No. of occurrence records = 120)
Model
SVM
RF
MAXENT
MARS
BRT
0.97
1
0.93
0.96
1
0.92
0.98
1
0.93
0.8
0.78
0.85
0.96
1
0.92
AUROC
Sensitivity
Specificity
Metric
Predicted probability
1
0.75
0.5
0.25
0

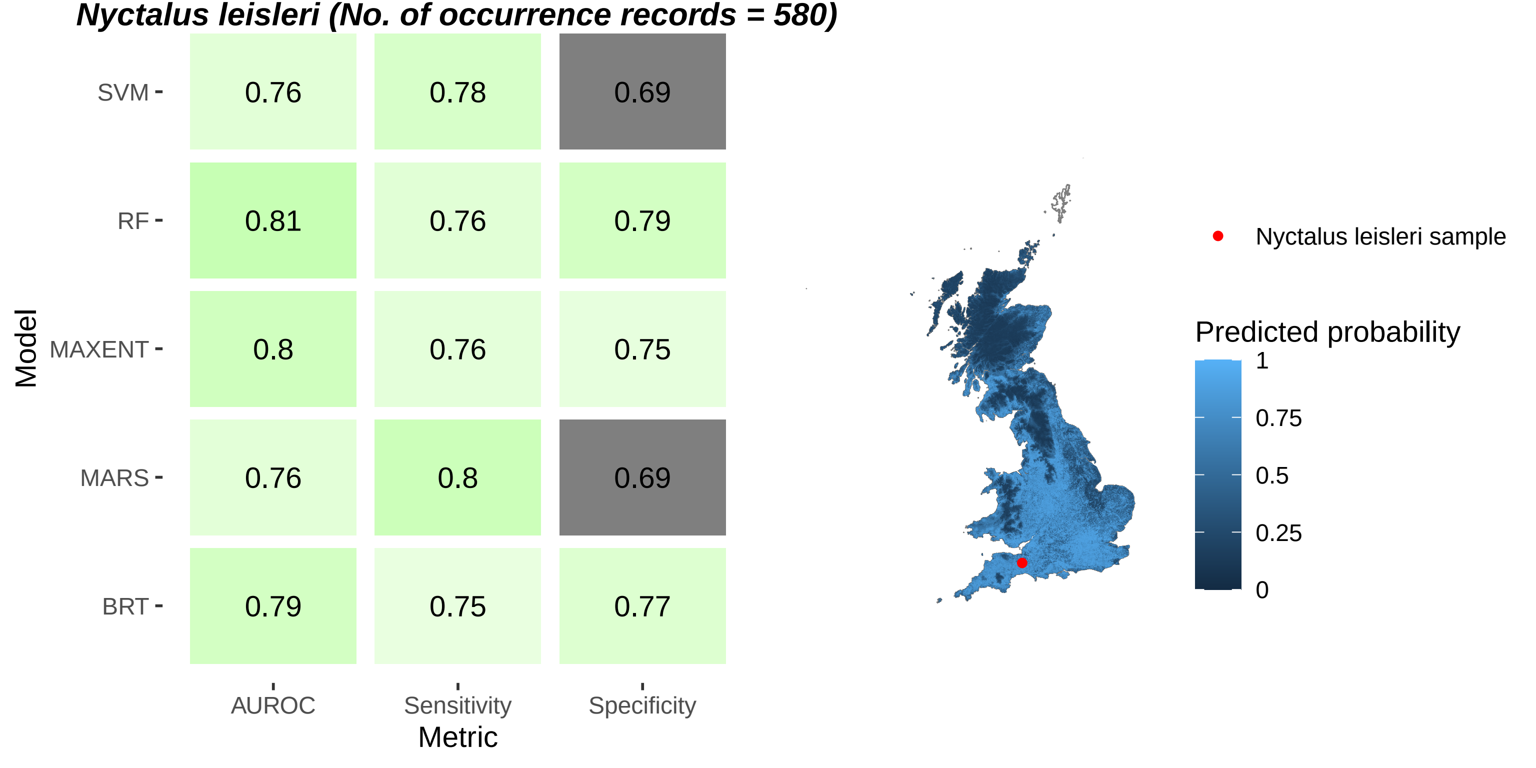
Nyctalus leisleri (No. of occurrence records = 580)
Model
SVM
RF
MAXENT
MARS
BRT
0.76
0.78
0.69
0.81
0.76
0.79
0.8
0.76
0.75
0.76
0.8
0.69
0.79
0.75
0.77
AUROC
Sensitivity
Specificity
Metric
Nyctalus leisleri sample
Predicted probability
1
0.75
0.5
0.25
0

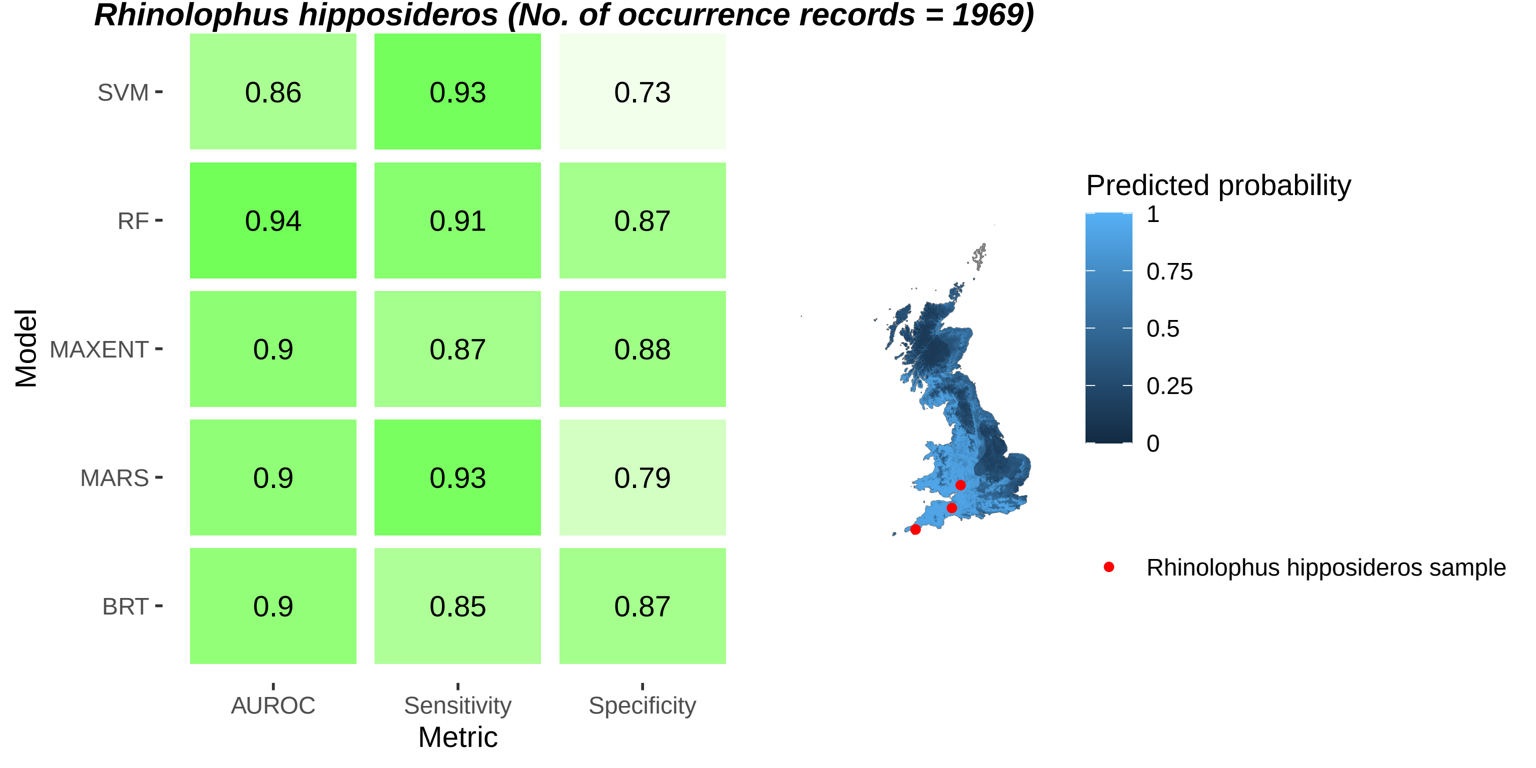
Rhinolophus hipposideros (No. of occurrence records = 1969)
Model
SVM
RF
MAXENT
MARS
BRT
AUROC
Sensitivity
Specificity
Metric
0.86
0.93
0.73
0.94
0.91
0.87
0.9
0.87
0.88
0.9
0.93
0.79
0.9
0.85
0.87
Predicted probability
1
0.75
0.5
0.25
0
Rhinolophus hipposideros sample

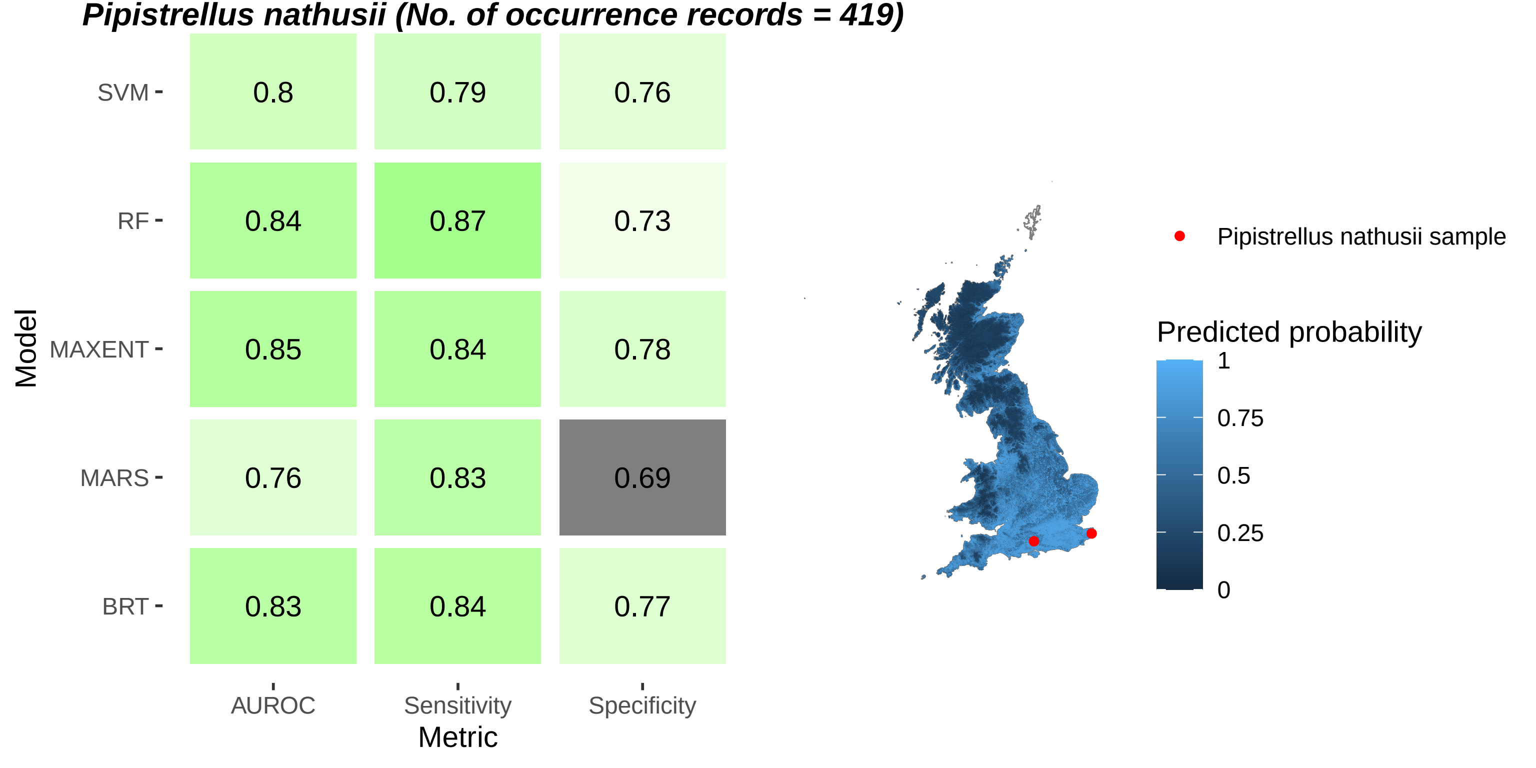
Pipistrellus nathusii (No. of occurrence records = 419)
SVM
RF
MAXENT
MARS
BRT
Model
0.8
0.79
0.76
0.84
0.87
0.73
0.85
0.84
0.78
0.76
0.83
0.69
0.83
0.84
0.77
AUROC
Sensitivity
Specificity
Metric
Pipistrellus nathusii sample
Predicted probability
1
0.75
0.5
0.25
0

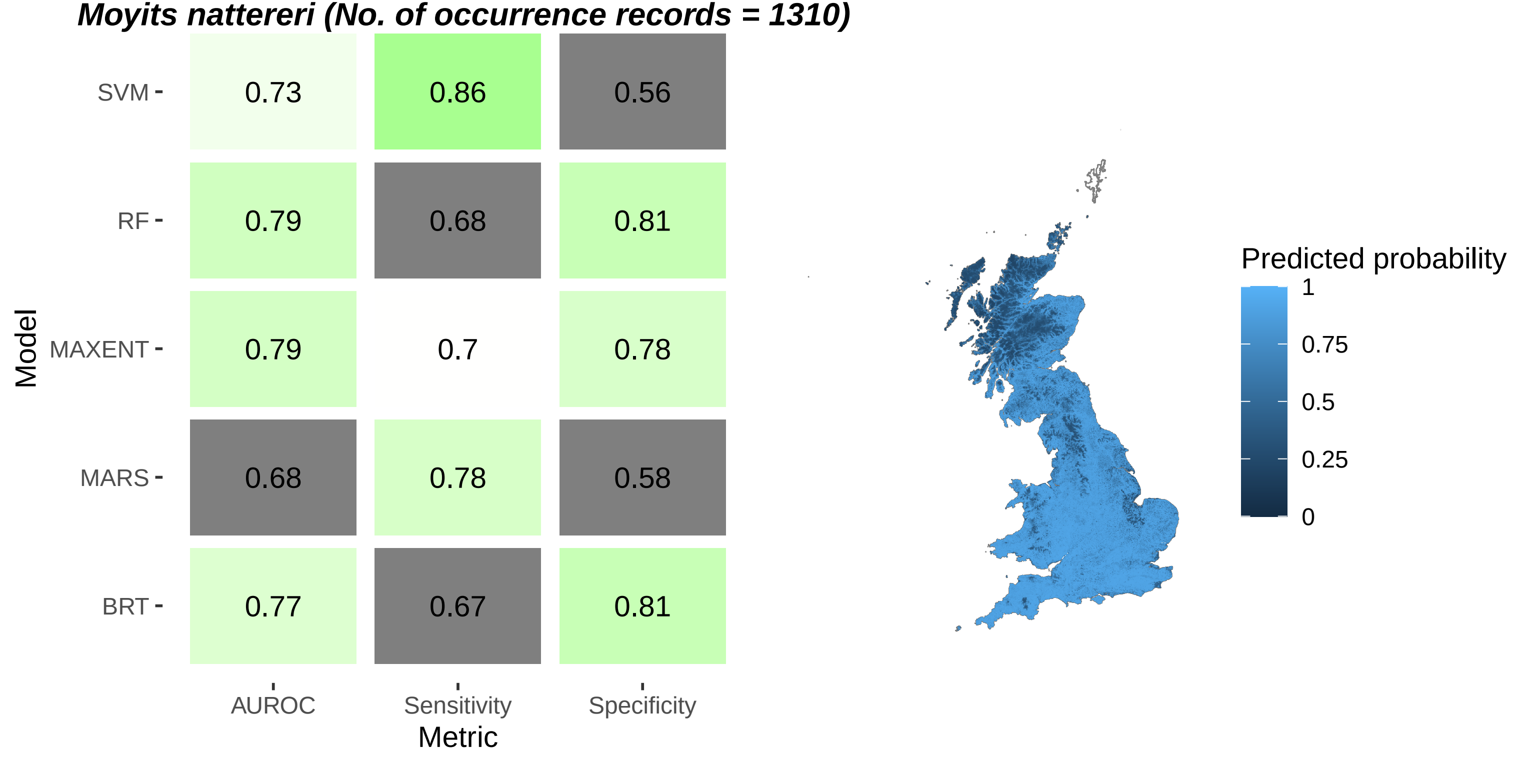
Moyits nattereri (No. of occurrence records = 1310)
Model
SVM
RF
MAXENT
MARS
BRT
0.73
0.86
0.56
0.79
0.68
0.81
0.79
0.7
0.78
0.68
0.78
0.58
0.77
0.67
0.81
AUROC
Sensitivity
Specificity
Metric
Predicted probability
1
0.75
0.5
0.25
0

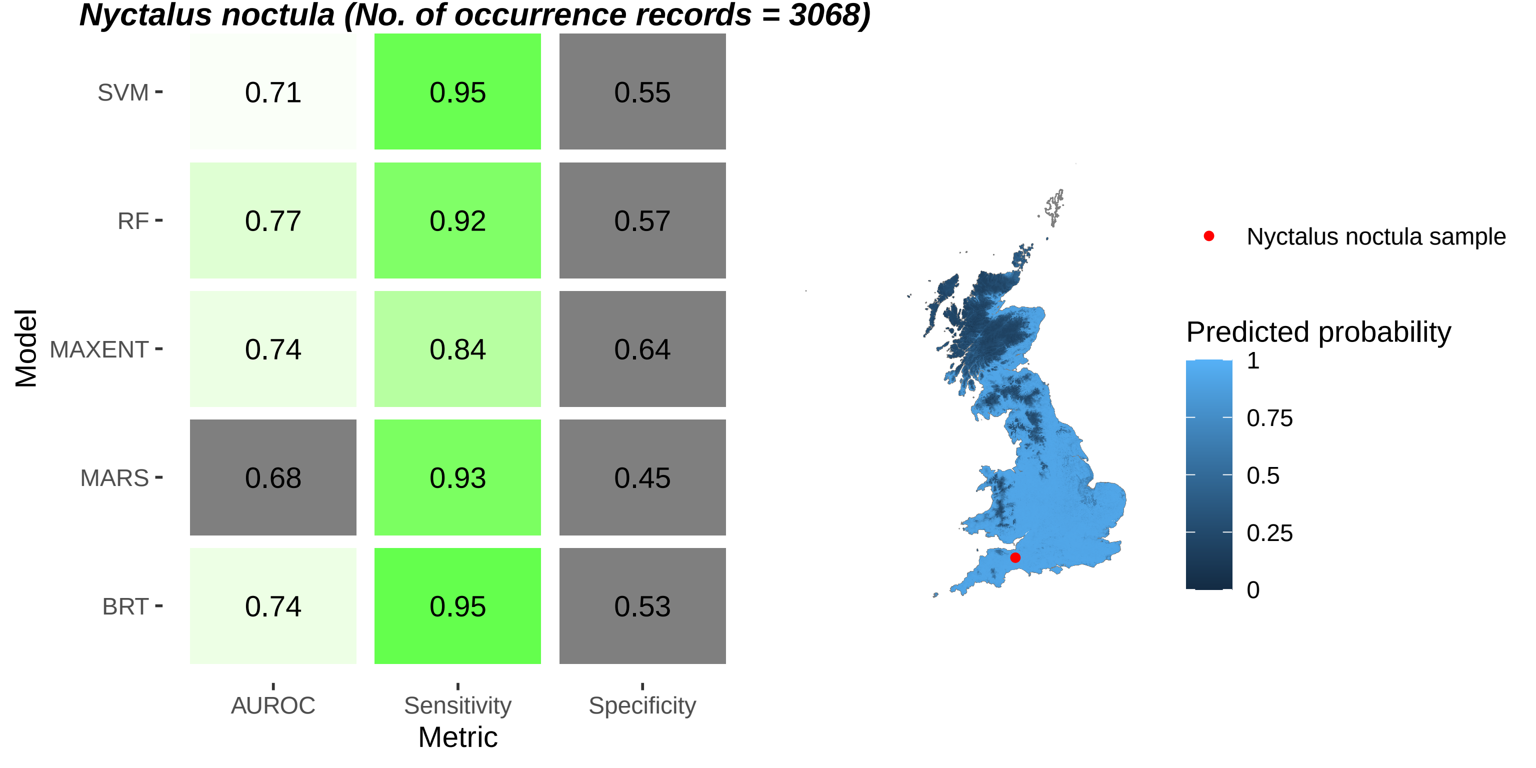
Nyctalus noctula (No. of occurrence records = 3068)
Model
SVM
RF
MAXENT
MARS
BRT
0.71
0.95
0.55
0.77
0.92
0.57
0.74
0.84
0.64
0.68
0.93
0.45
0.74
0.95
0.53
AUROC
Sensitivity
Specificity
Metric
Nyctalus noctula sample
Predicted probability
1
0.75
0.5
0.25
0

***Eptesicus serotinus* (No. of occurrence records = 1702)**

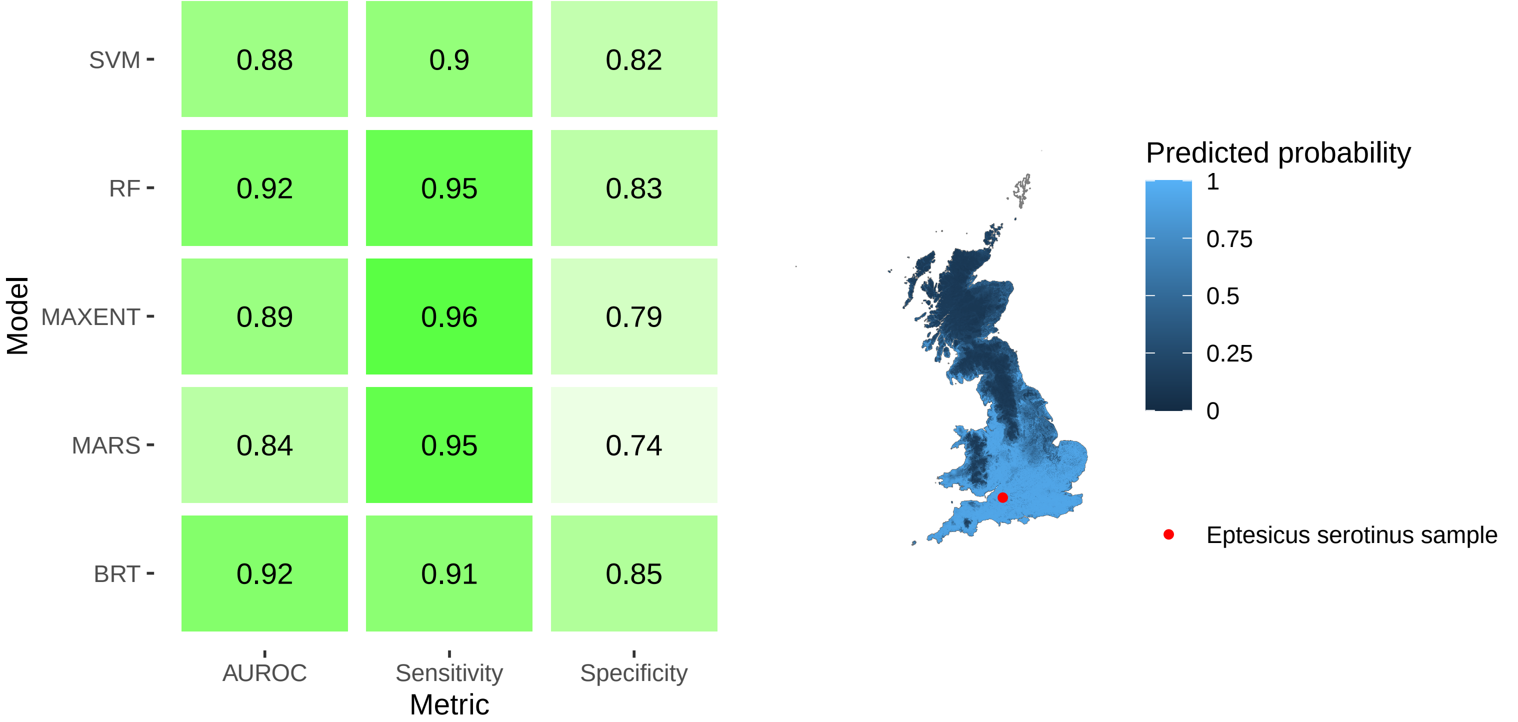

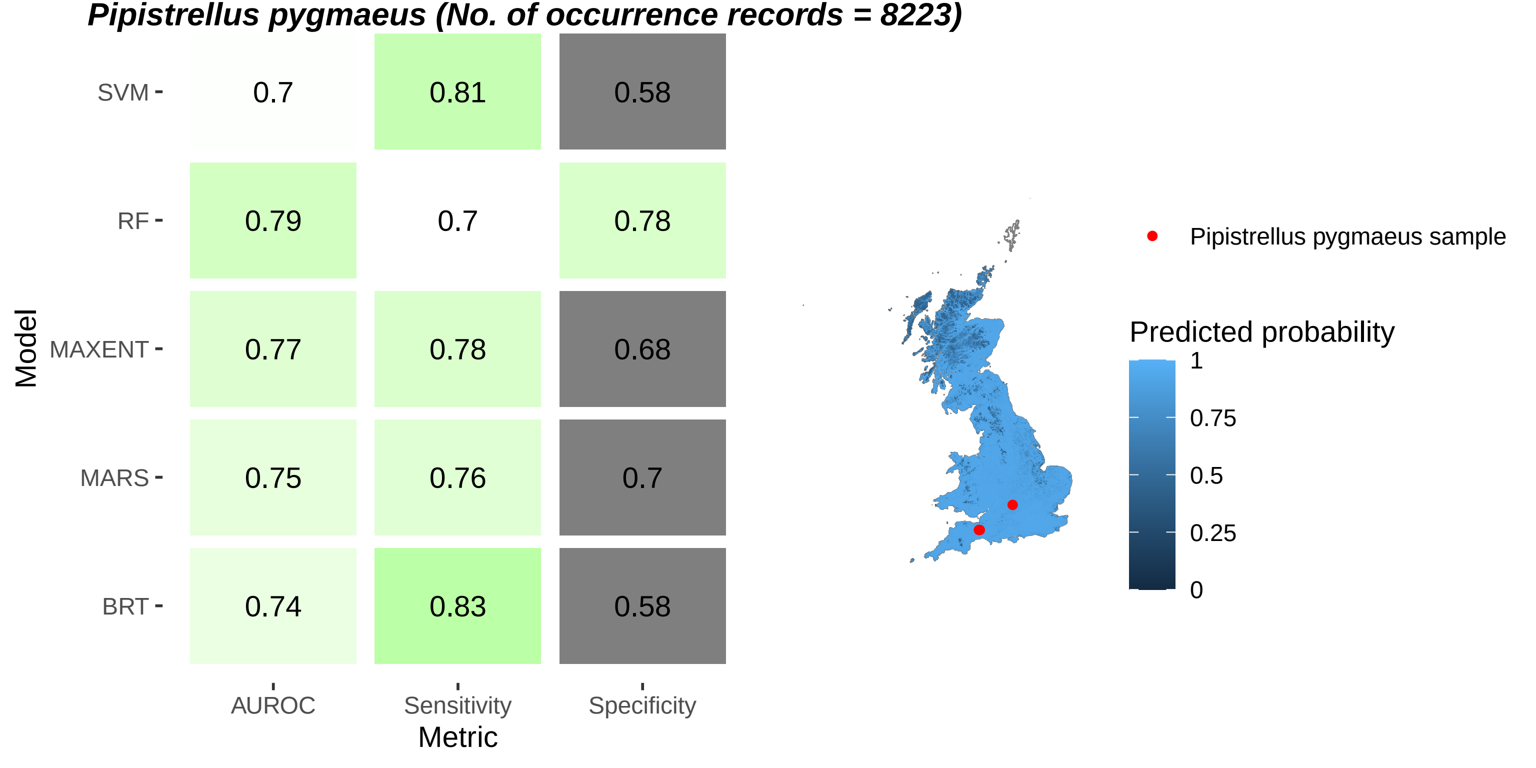
Pipistrellus pygmaeus (No. of occurrence records = 8223)
Model
SVM
RF
MAXENT
MARS
BRT
0.7
0.81
0.58
0.79
0.7
0.78
0.77
0.78
0.68
0.75
0.76
0.7
0.74
0.83
0.58
AUROC
Sensitivity
Specificity
Metric
Pipistrellus pygmaeus sample
Predicted probability
1
0.75
0.5
0.25
0

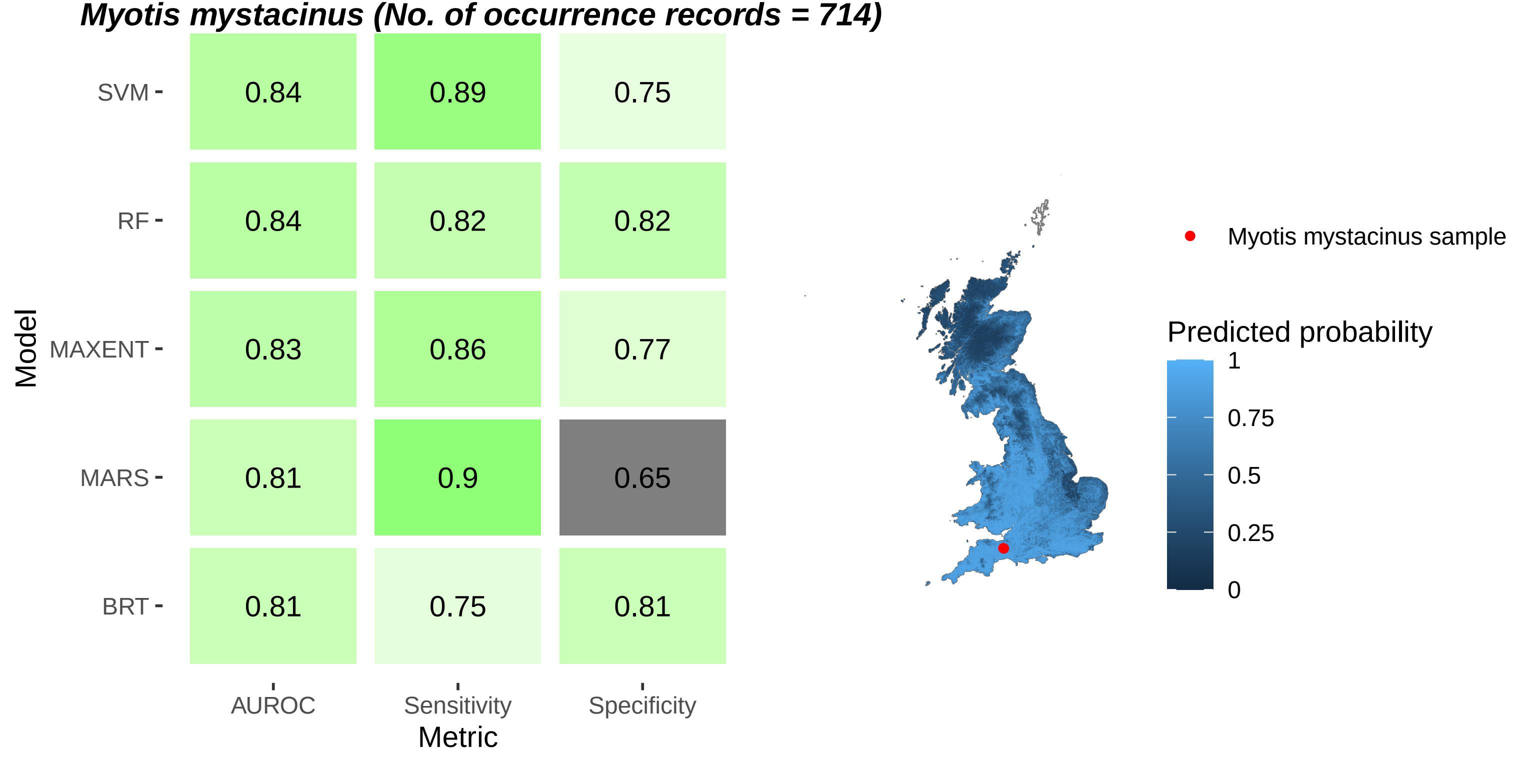
Myotis mystacinus (No. of occurrence records = 714)
SVM
RF
MAXENT
MARS
BRT
Model
0.84
0.89
0.75
0.84
0.82
0.82
0.83
0.86
0.77
0.81
0.9
0.65
0.81
0.75
0.81
AUROC
Sensitivity
Specificity
Metric
Myotis mystacinus sample
Predicted probability
1
0.75
0.5
0.25
0
