## Supplementary figures and images for "Genomic screening of 16 UK native bat species through conservationist networks uncovers coronaviruses with zoonotic potential"

### Supplementary Figure 1.pdf

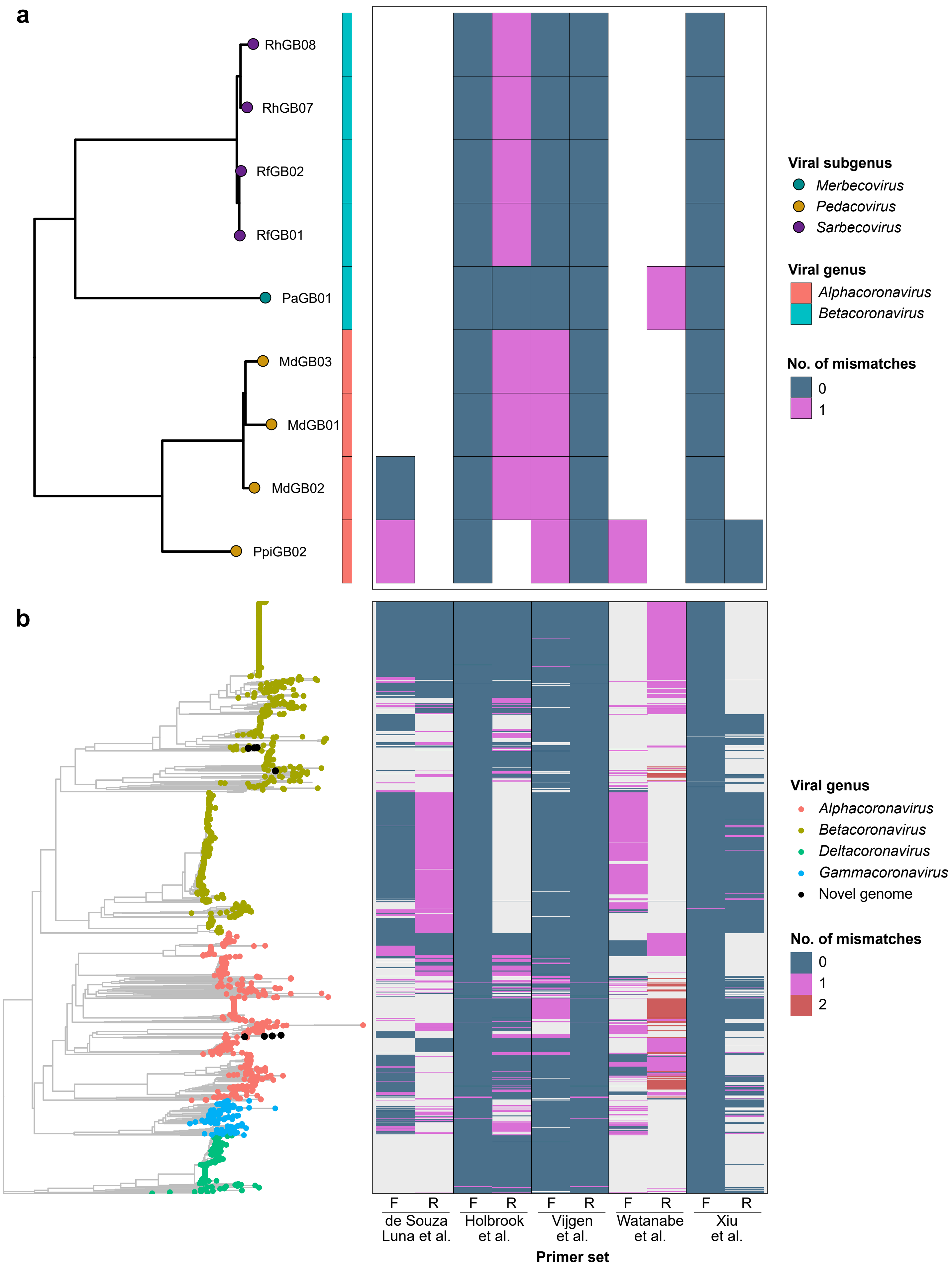

### Supplementary Figure 2.pdf

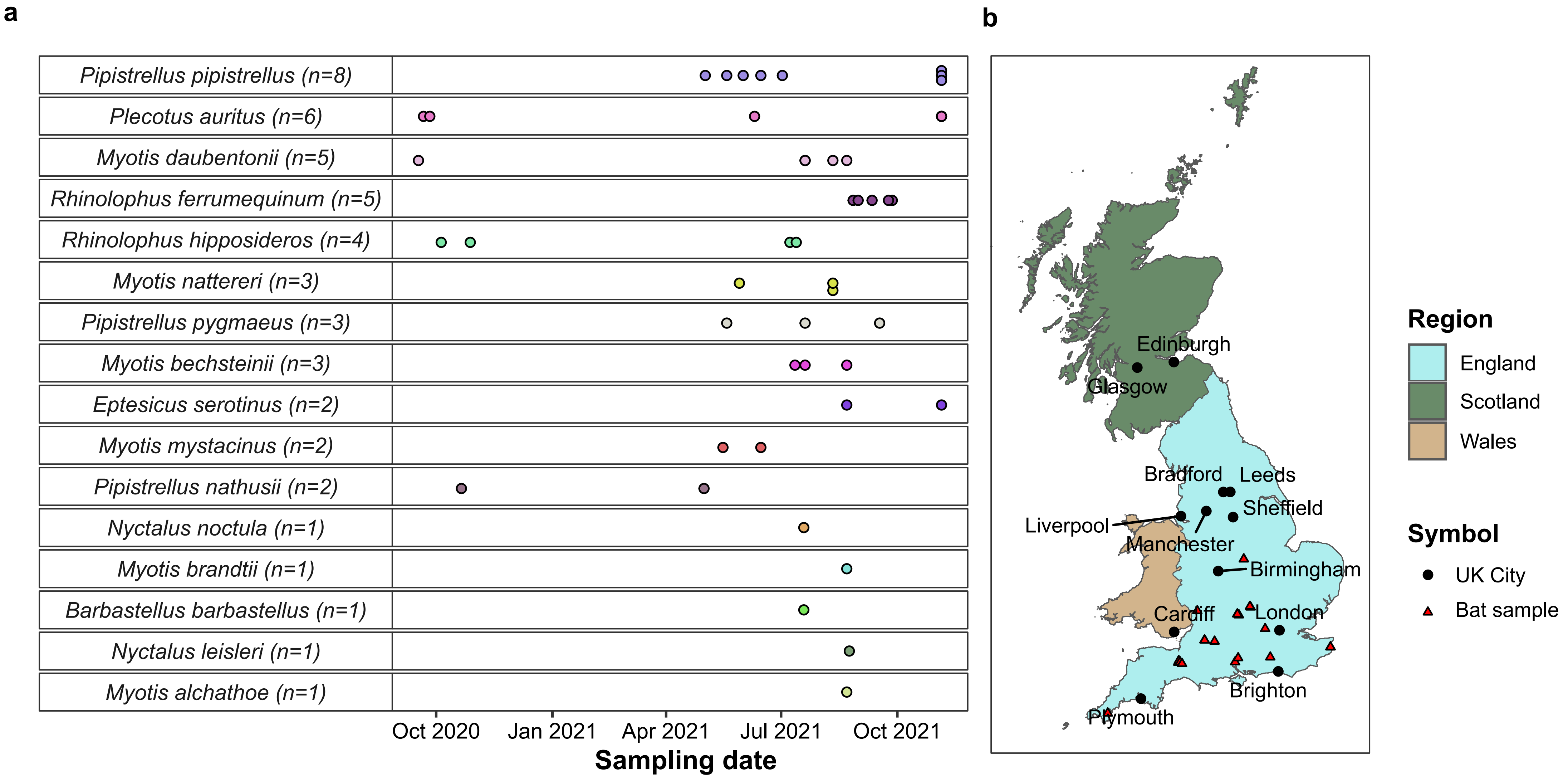

### Supplementary Figure 3.pdf

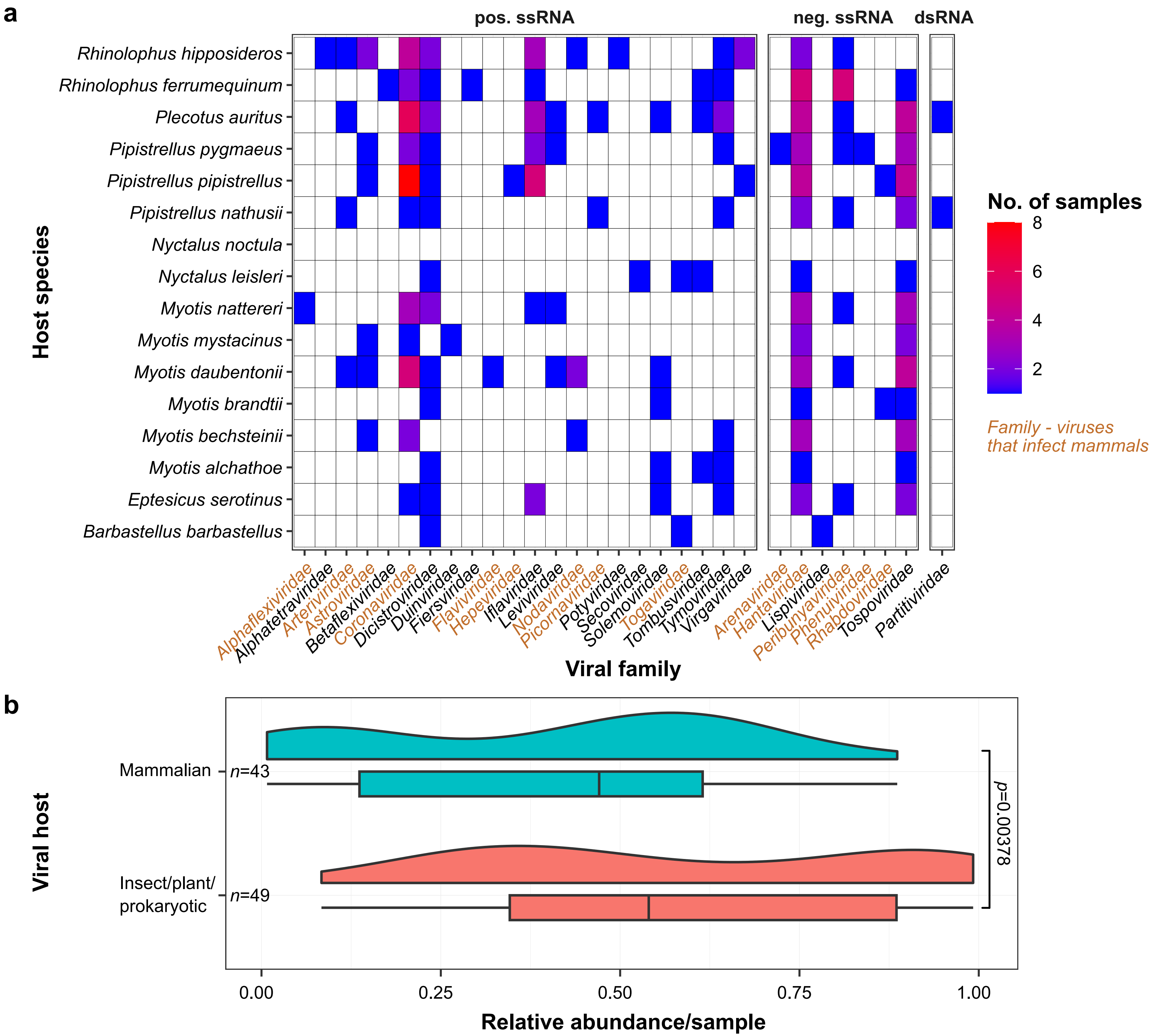

### Supplementary Figure 4.pdf

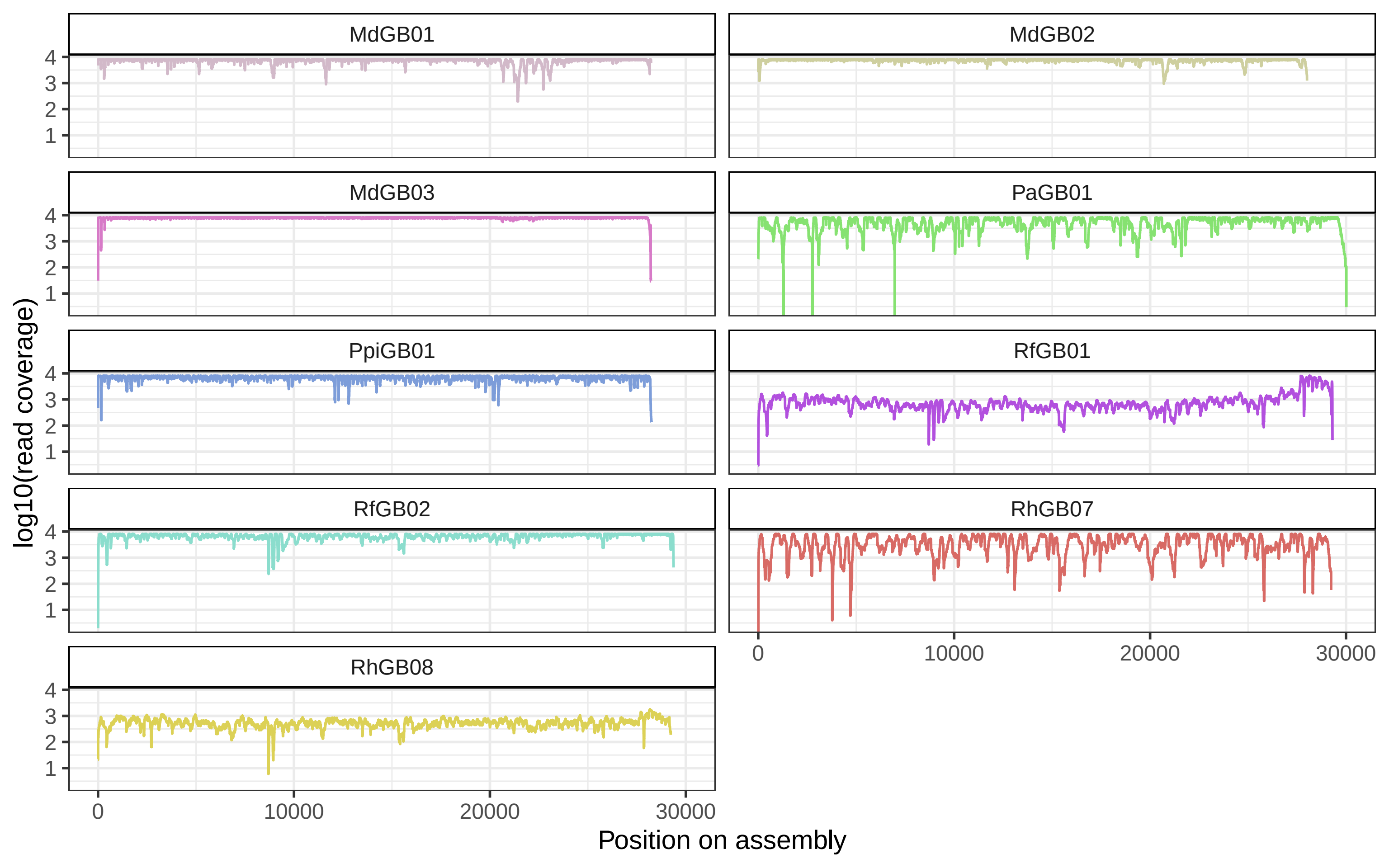

### Supplementary Figure 5.pdf

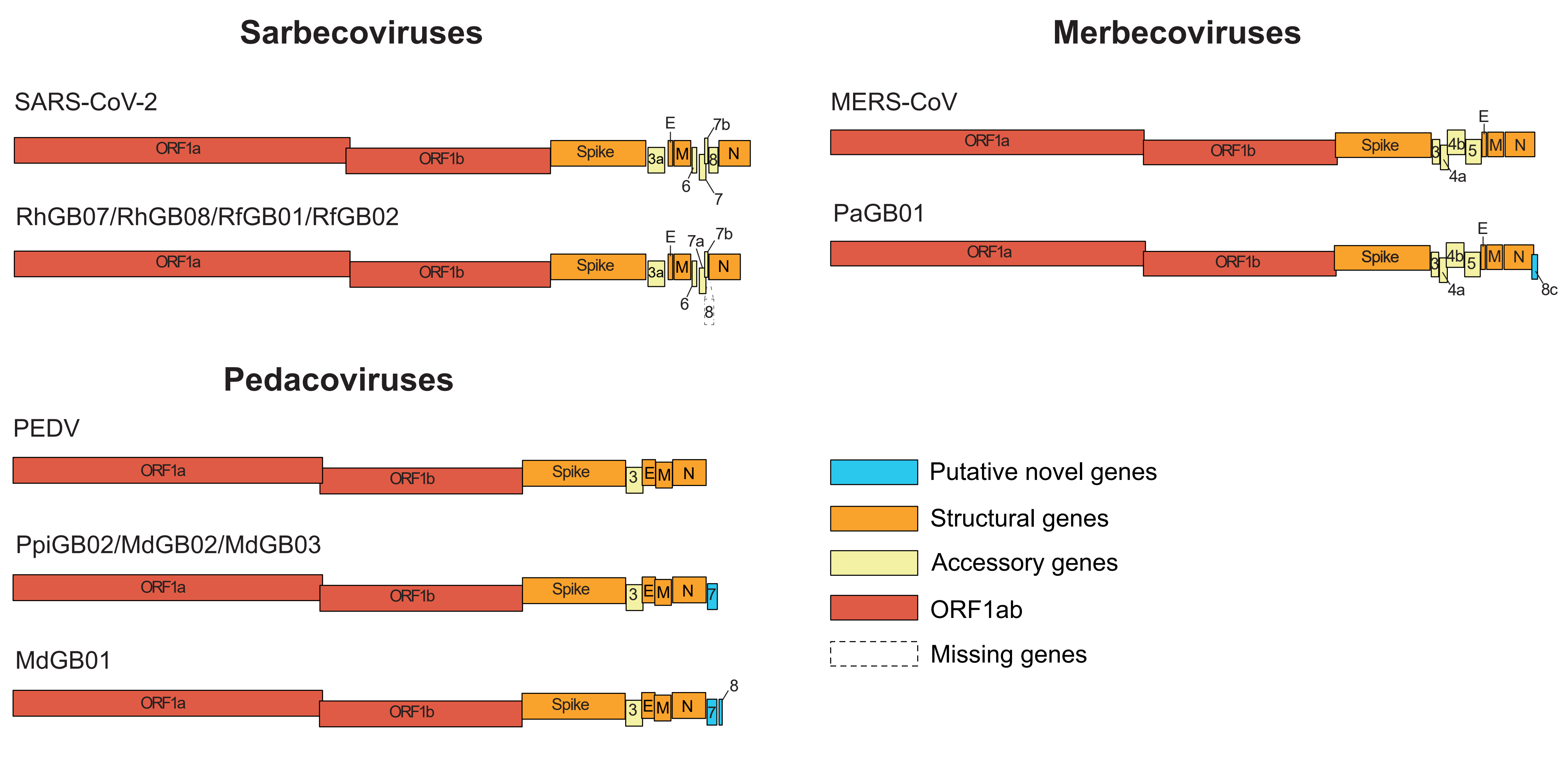

### Supplementary Figure 6.pdf

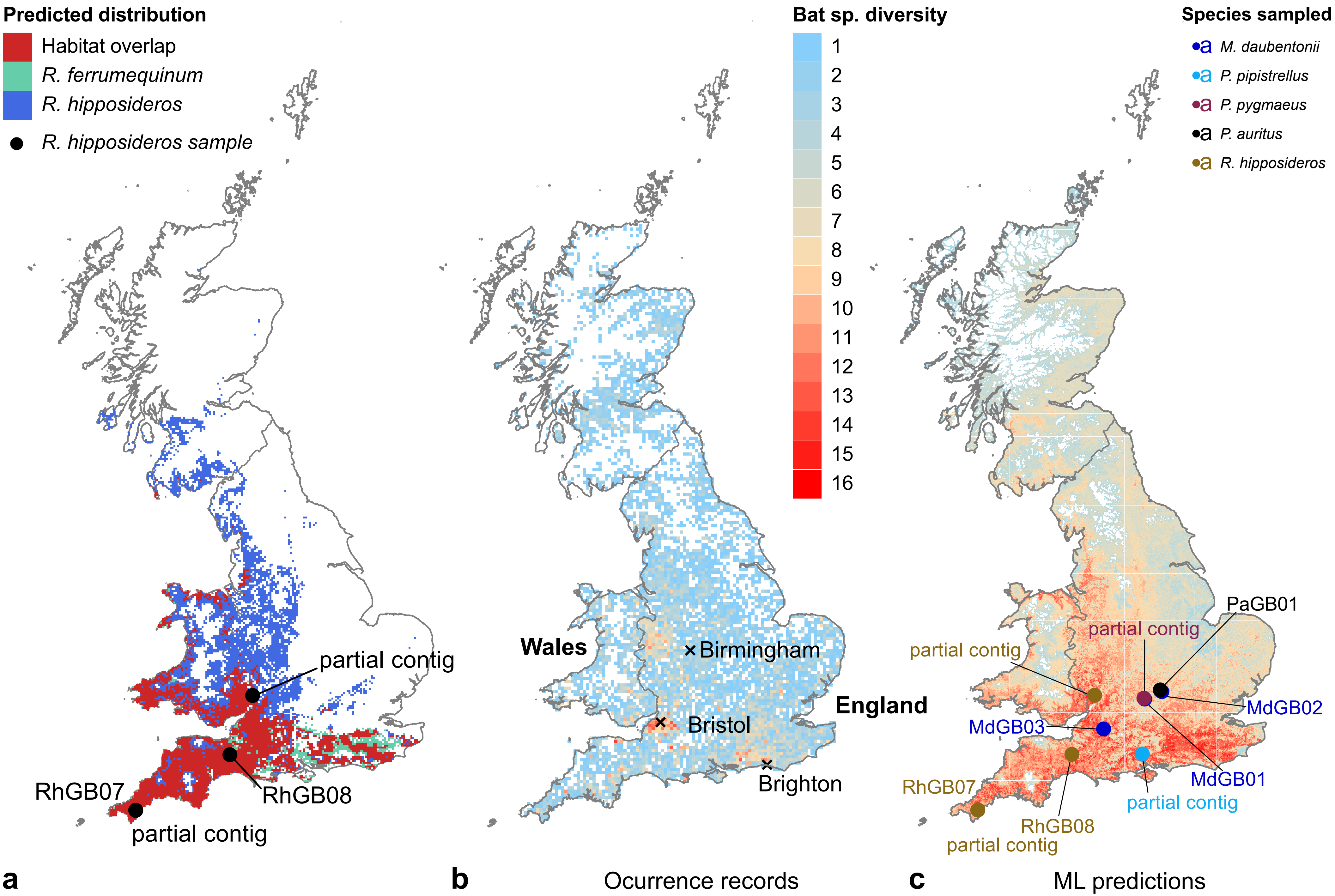

### Supplementary Figure 7.pdf

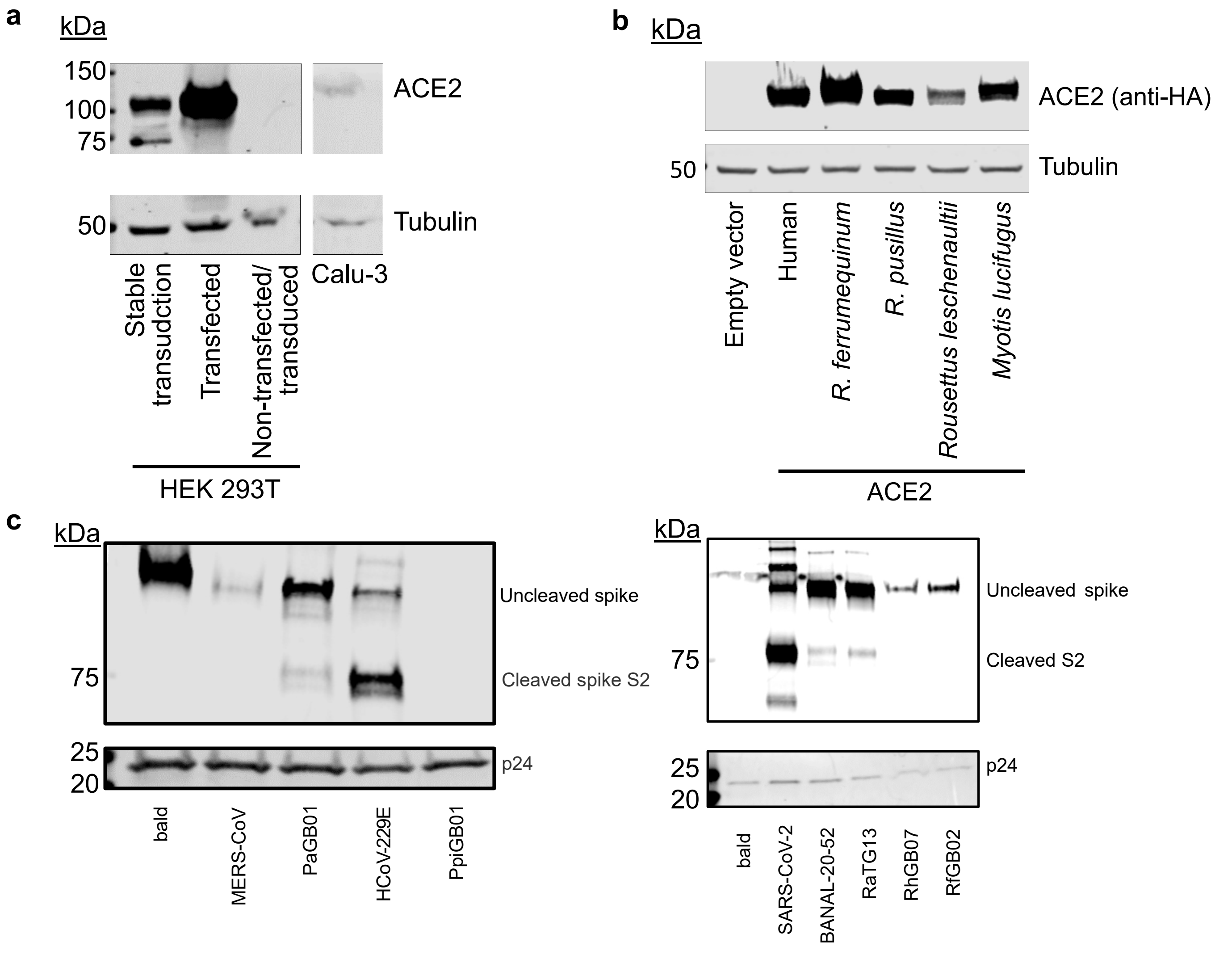

### Supplementary Figure 8.pdf

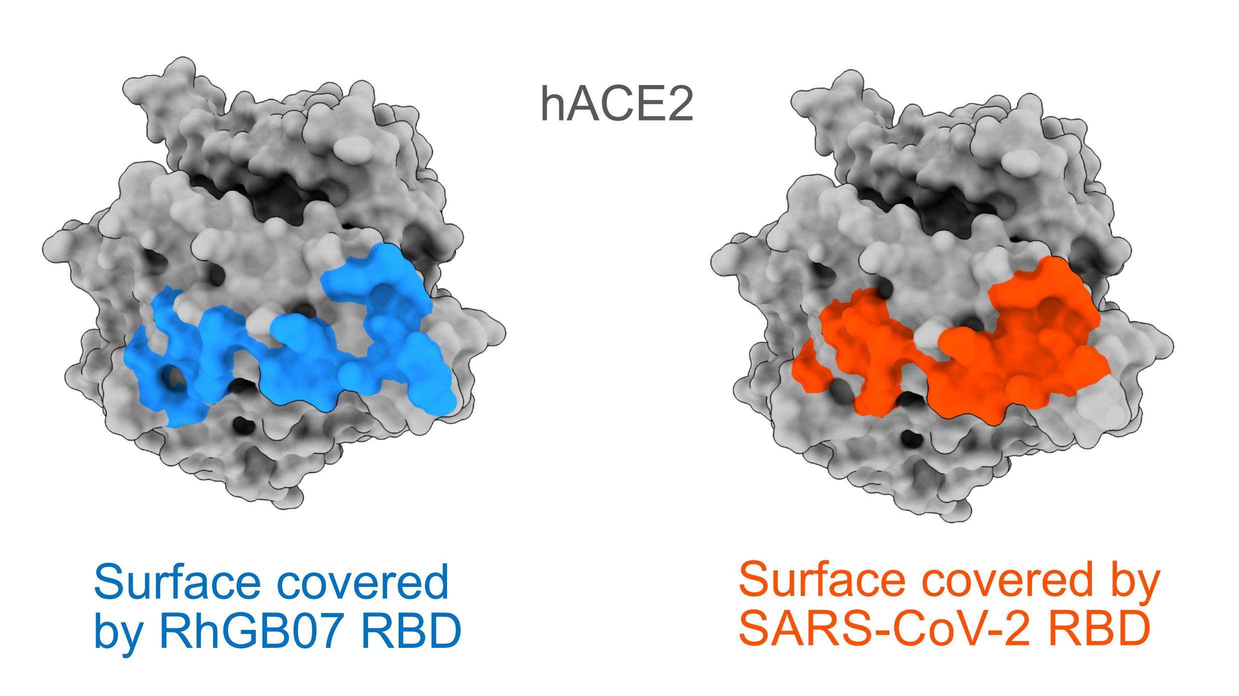

### Supplementary Figure 9.pdf

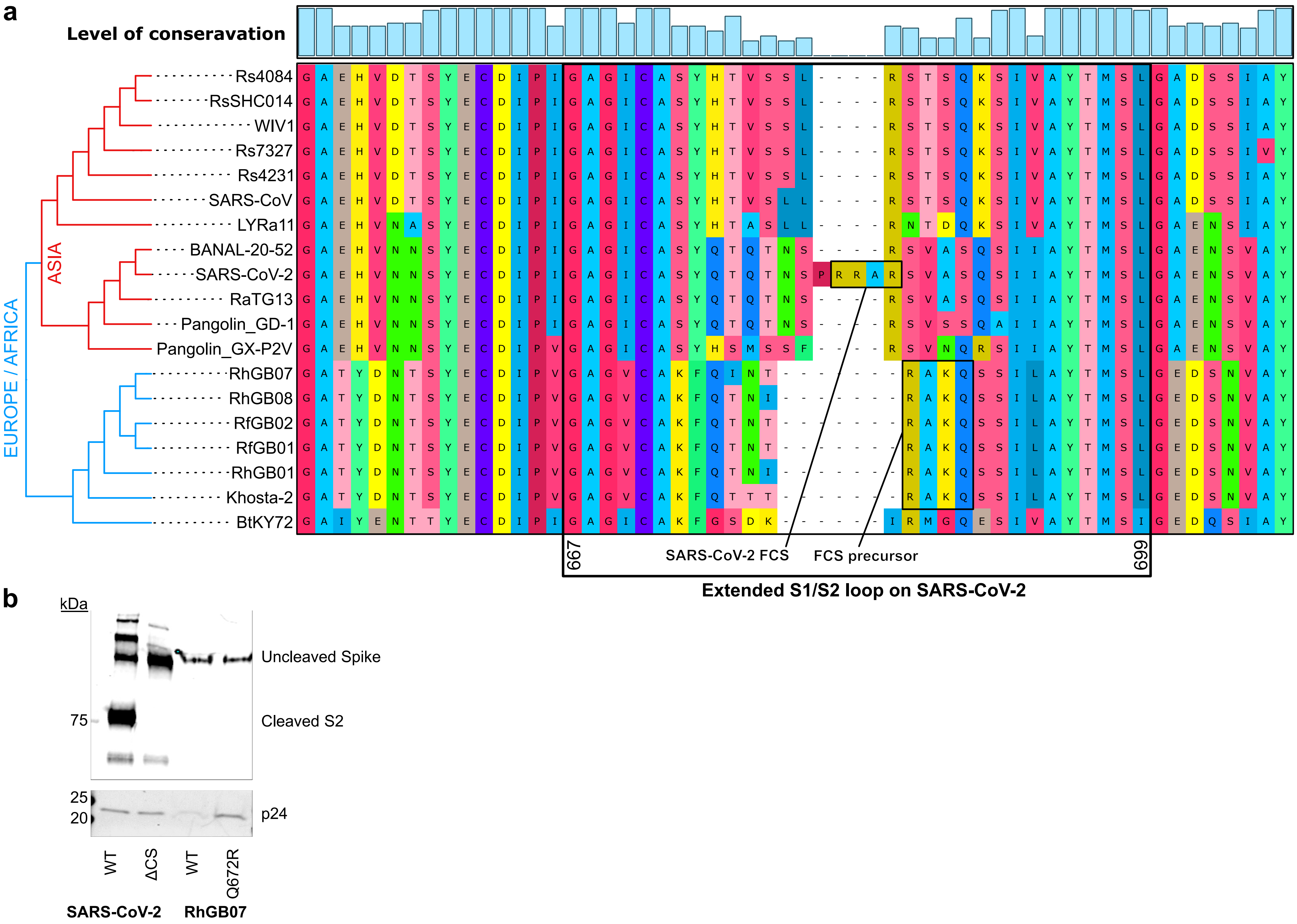

### Supplementary Figure 10.pdf

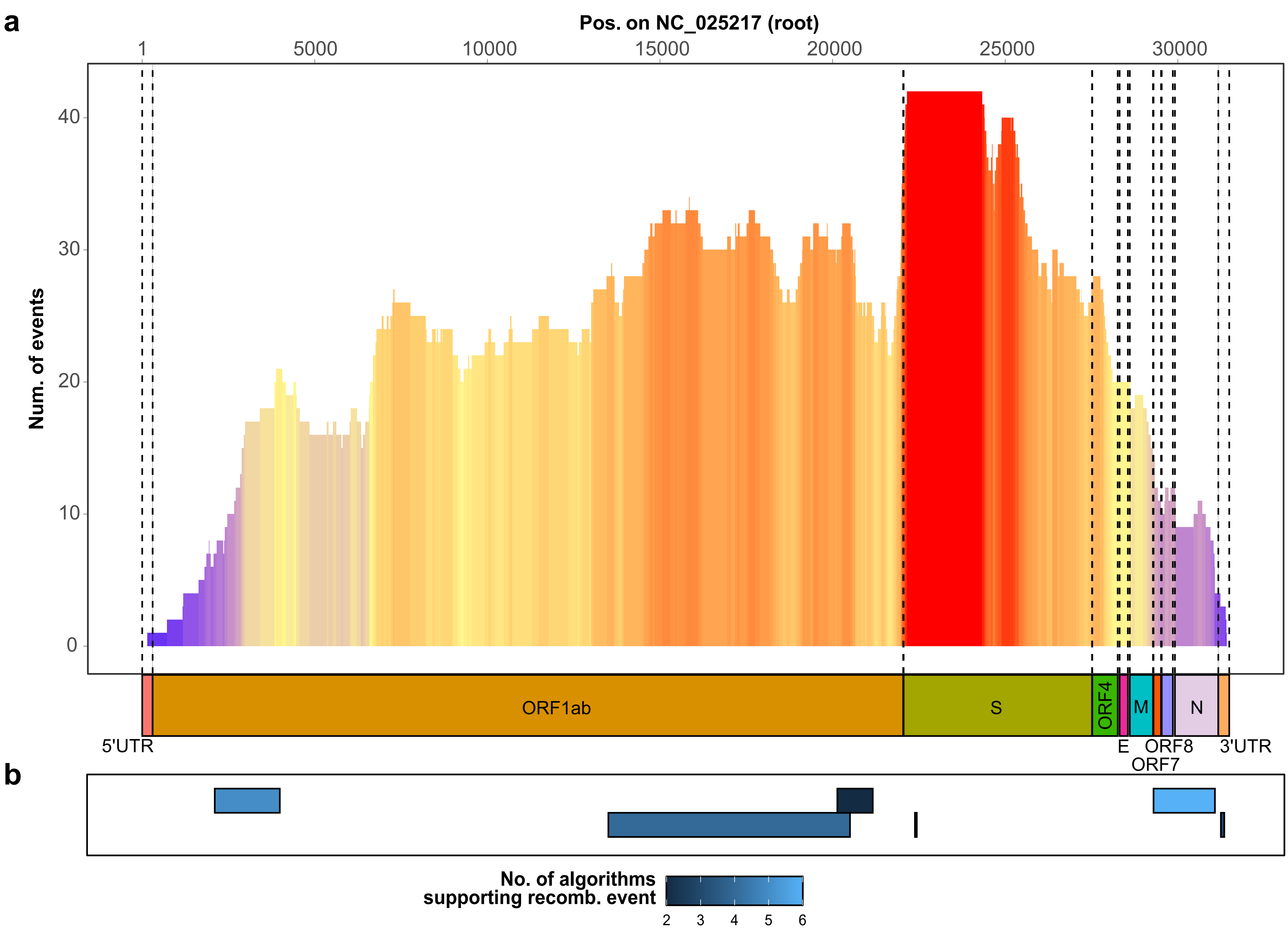

### Supplementary Figure 12.pdf

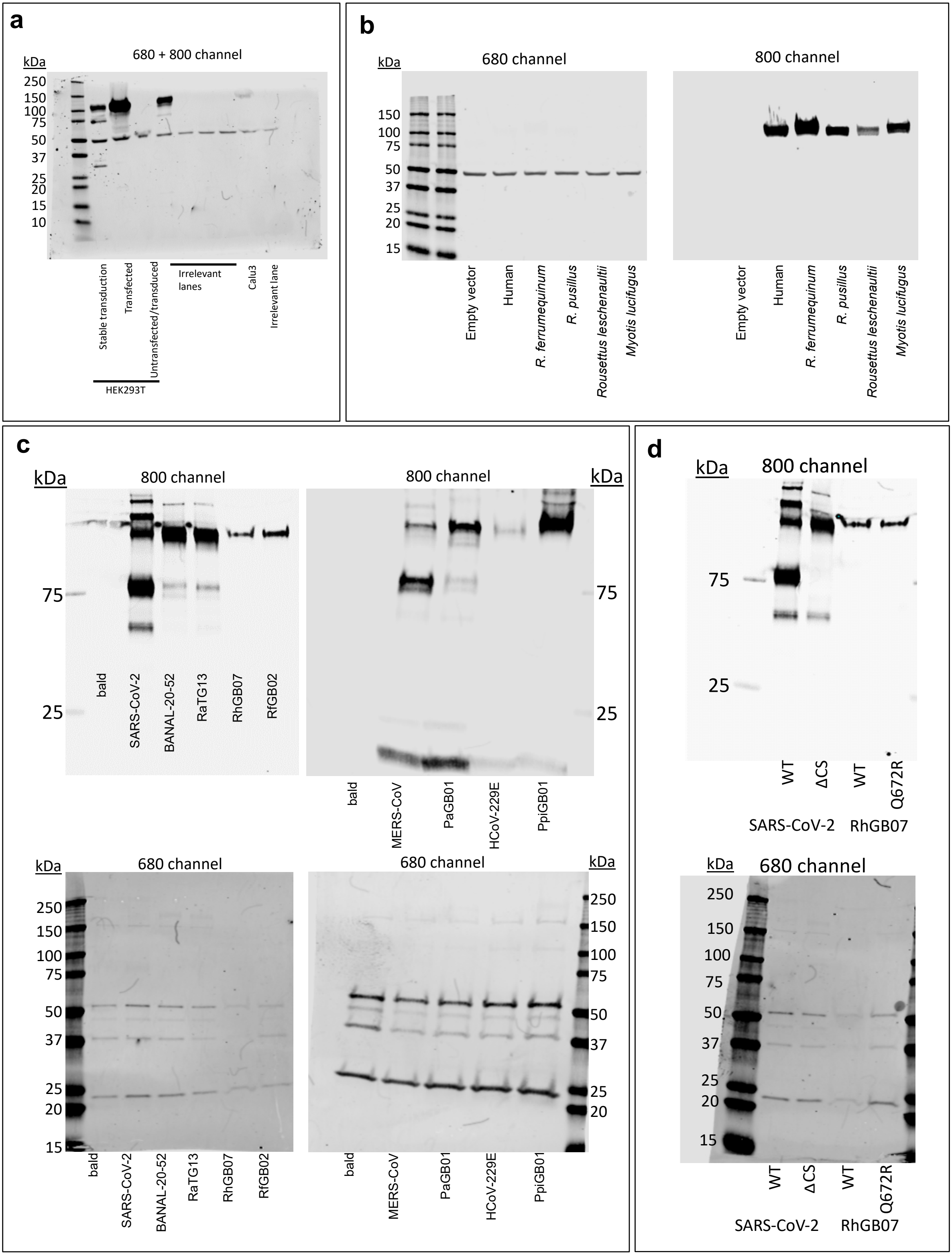
